# Non-selective beta-blockers reduce bystander CD8^+^ T cell activation in decompensated liver cirrhosis

**DOI:** 10.1101/2025.11.18.688703

**Authors:** Ayesha Lietzau, Ines Tapken, So-Young Kim, Roni Souleiman, Erich Freyer, Anke RM Kraft, Sarah Schütte, Tammo L Tergast, Benjamin Maasoumy, Heiner Wedemeyer, Isabell Drath, Gabriele Zurek, Eui-Cheol Shin, Markus Cornberg, Christian Niehaus

**Affiliations:** Department of Gastroenterology, Hepatology, Infectious Diseases and Endocrinology, Hannover Medical School, Hannover, Germany; Centre for Individualised Infection Medicine (CiiM), a joint venture between the Helmholtz Centre for Infection Research (HZI) and Hannover Medical School (MHH), Hannover, Germany; Twincore, Centre for Experimental and Clinical Infection Research, a joint venture between the Helmholtz Centre for Infection Research (HZI) and the Hannover Medical School, Hannover, Germany; Laboratory of Immunology and Infectious Diseases, Graduate School of Medical Science and Engineering, Korea Advanced Institute of Science and Technology (KAIST), Daejeon 34051, Republic of Korea; German Center for Infection Research (DZIF), Partner-site Hannover-Braunschweig, Hannover, Germany; Cluster of Excellence RESIST (EXC 2155), Hannover Medical School, Hannover, Germany; German Center for Infection Research, HepNet Study-House German Liver Foundation, Hannover, Germany; Department of Pharmacology, Toxicology and Pharmacy, University of Veterinary Medicine Hannover, Germany; Medizinisches Labor Bremen, Bremen, Germany

**Keywords:** Bystander activation, cirrhosis-associated immune dysfunction, liver cirrhosis, NSBB, systemic inflammation, T cells

## Abstract

Cirrhosis is characterized by immune dysfunction in which activated CD8⁺ T cells fuel systemic inflammation and disease progression. Non-selective beta-blockers (NSBB), widely prescribed for portal hypertension, have incompletely understood immunomodulatory effects. Here we show that CD8⁺ T cells express β2-adrenergic receptors, enriched in bystander relative to antigen-specific subsets. *In vitro*, the NSBB propranolol selectively suppressed cytokine-driven bystander activation, reducing CD69⁺CXCR6⁺ and NKG2D⁺ populations and pro-inflammatory cytokines, while preserving antigen-specific responses. Transcriptomic profiling after NSBB treatment revealed downregulation of interferon signaling pathway via STAT1. In paired blood and ascites samples from patients with decompensated cirrhosis (n = 31), NSBB therapy was associated with reduced bystander-activated CD8⁺ T cells. In a retrospective cohort (n = 624), NSBB therapy correlated with lower systemic and intrahepatic inflammation. These findings identify β-adrenergic blockade as a mechanism that restrains bystander CD8⁺ T cell responses without impairing antigen-specific immunity, supporting NSBB therapy as a strategy to mitigate inflammation and improve outcomes in cirrhosis.

**Graphical abstract:** 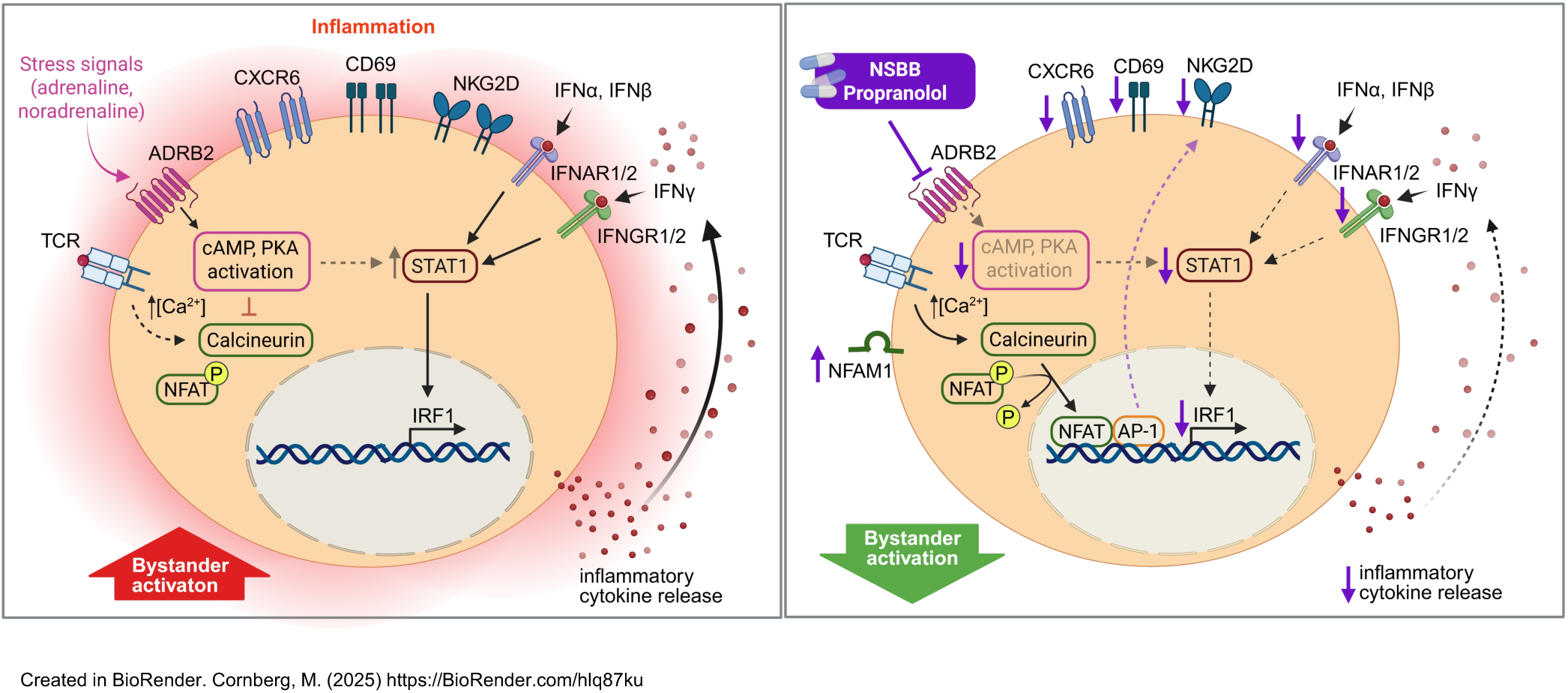

## Introduction

Liver cirrhosis is a leading cause of morbidity and mortality worldwide, accounting for more than one million deaths annually (1, 2). The most common causes of liver cirrhosis are chronic hepatitis virus infection, such as hepatitis B and hepatitis C, alcohol-related liver disease, and metabolic dysfunction-associated steatotic liver disease (2). Disease progression is driven not only by declining liver function and portal hypertension, leading to clinical complications such as decompensation (1), but also by cirrhosis-associated immune dysfunction (CAID), a state of combined immunodeficiency and systemic inflammation (3). In advanced stages of cirrhosis, this inflammatory response becomes the major driver of acute-on-chronic liver failure (ACLF) and subsequently poor clinical outcomes (4–7).

Among immune cells, bystander-activated CD8⁺ T cells have emerged as critical amplifiers of inflammation in advanced liver disease (8–12). Unlike antigen-specific CD8⁺ T cells, which are activated through the T cell receptor (13), these cells are triggered by pro-inflammatory cytokines such as IL-12, IL-18 (11, 14) and IL-15 (11, 15), and have been shown to drive liver pathology through non-specific cytotoxicity in MASH (8) and chronic hepatitis virus infection (9, 12). Extending these findings, we recently identified an enrichment of bystander-activated CD8^+^ T cells co-expressing CD69 and CXCR6 in the ascites of patients with decompensated liver cirrhosis, which correlate with disease severity (11). Yet, therapeutic strategies directly targeting this pathogenic T cell subset are lacking.

Non-selective beta-blockers (NSBB) are established as the standard of care in cirrhosis, where they reduce portal hypertension and thereby lower the risk of variceal bleeding and decompensation (16–18). Intriguingly, NSBB therapy has also been consistently associated with improved survival and reduced systemic inflammation in clinical studies (18–23). These anti-inflammatory effects have been largely attributed to reduced intestinal permeability and bacterial translocation (19, 21, 23). While previous studies have reported direct effects of NSBB on lymphocytes via beta-adrenergic receptors (ADRB) (24–28), including CD8^+^ T cells (29–32), their immunomodulatory impact on bystander CD8^+^ T cells in cirrhosis remains unclear.

Here, we identify a novel mechanism by which the NSBB propranolol directly modulates the immune system in cirrhosis. Using *in vitro* functional assays, high-dimensional phenotyping and transcriptomic profiling we show that propranolol selectively suppresses bystander-activated CD8⁺ T cells by targeting the STAT1/IRF1 axis, while preserving antigen-specific immunity. These findings expand the therapeutic scope of NSBB, highlighting their potential as immunomodulatory agents to attenuate inflammation in cirrhosis.

## Results

### ADRB2 is enriched in effector-memory CD8⁺ T cells and marks bystander over antigen-specific CD8⁺ T cells

We first investigated whether CD8⁺ T cells from patients with decompensated cirrhosis express beta-adrenergic receptors as potential targets of NSBB therapy. Flow cytometric staining of freshly isolated PBMC and MNC revealed surface expression of ADRB1 and, more predominantly, ADRB2 in blood and ascites CD8^+^ T cells (Fig.1a, b). When comparing matched blood to ascites CD8^+^ T cells, no significant differences in ADRB1 and ADRB2 expression were observed (Extended Data Fig. 2a). In line with this, analysis of a publicly available single-cell RNA-sequencing dataset (GSE182159) (33) of matched liver- and blood-derived CD8⁺ T cells revealed that ADRB2, but not ADRB1, was present at significant levels on liver CD8^+^ T cells (Extended Data Fig. 2b-f). Next, we re-analyzed previously published single-cell RNA-sequencing data obtained from patients with acute HAV infection (15), where the bystander activation of memory CD8^+^ T cells is well-exemplified (34). Peripheral blood CD8^+^ T cells were sorted from patients with acute HAV infection (n = 5) and healthy donors (n = 3) and stained with DNA-barcoded MHC-I multimers (dCODE dextramers) to detect CD8^+^ T cells specific for HAV, HCMV, EBV, and influenza A virus (15). CD8^+^ T cells were further sorted into dCODE dextramer-positive and -negative cells, and two sorted cell populations were re-mixed at a 1:9 ratio. Re-mixed cells were further labeled with antibody-derived tags (ADTs), and scRNA-seq was performed (Fig. 1c) (15). We first confirmed that CD8⁺ T cells almost exclusively expressed ADRB2, whereas ADRB1 and ADRB3 were nearly absent (Fig. 1d). ADRB2⁺ CD8⁺ T cells were found within the effector/memory cluster (Fig. 1e), and the increased expression of ADRB2 in the effector/memory cluster compared to the naïve cluster was confirmed (Fig. 1f). Further clustering of effector/memory CD8⁺ T cells into eight subclusters (C0–C7; Fig. 1g) revealed ADRB2 enrichment in the C3 subcluster, corresponding to senescent effector-memory CD8⁺ T cells (Fig. 1h, i), characterized by the upregulation of CD57, loss of CD28 expression, and a pro-inflammatory profile with increased IFNG and TNF production (Fig. 1j, k). Given that HAV infection triggers a robust immune response with massive cytokine release, activating both bystander and antigen-specific CD8^+^ T cells (34, 35), we used this virus model to compare ADRB2 expression levels between these two populations. Notably, ADRB2 expression was higher in bystander-activated CD8^+^ T cells, defined as HAV-non-specific T cells that respond to heterologous viral antigens (HCMV, EBV, IAV), than in HAV-specific CD8^+^ T cells, within patients with acute HAV infection (Fig. 1l, m). Similar levels were observed in bystander CD8^+^ T cells from patients with acute HAV infection and healthy donors (Fig. 1l, m).

**Fig. 1.**
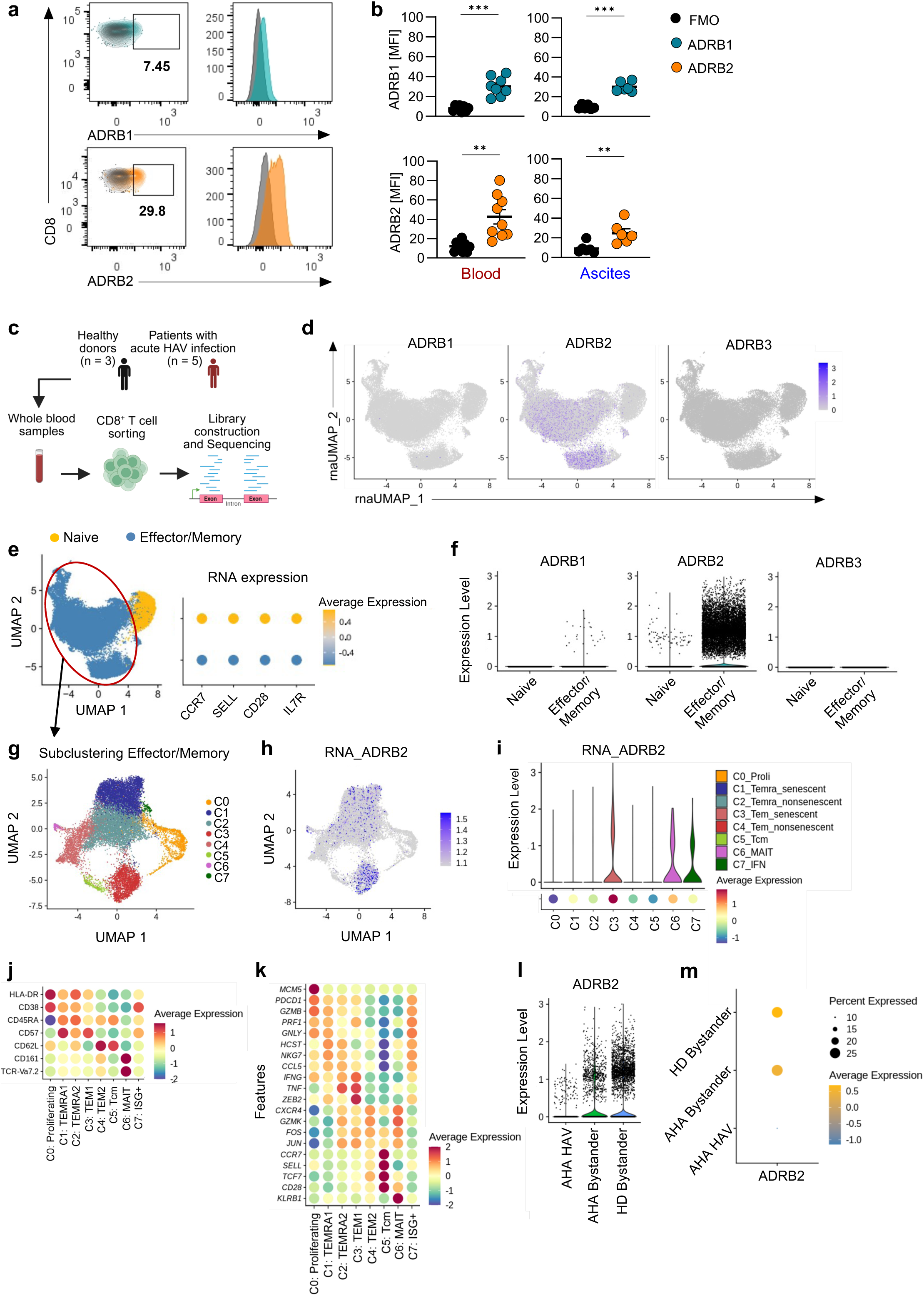
ADRB2 is enriched in effector-memory CD8⁺ T cells and marks bystander over antigen-specific CD8⁺ T cells. (a) Representative FACS plots and histograms displaying ADRB1 and ADRB2 expression on CD8^+^ T cells in the blood of a patient with decompensated cirrhosis. (b) Expression of ADRB1 and ADRB2 on CD8^+^ T cells in blood (n = 8-9) and ascites (n = 6) from patients with decompensated cirrhosis, shown as mean fluorescence intensity (MFI) versus FMO control. Paired *t* test. (c) Schematic workflow. Previously published scRNA-seq data from HAV-infected patients (n = 5) and healthy donors (n = 3) were reanalyzed (15). CD8⁺ T cells were isolated and labeled with DNA-barcoded MHC-I multimers (dCODE dextramers) to distinguish HAV-specific CD8^+^ T cells from those specific for HCMV, EBV and IAV, and analyzed using CITE-seq. Created in BioRender. Cornberg, M. (2025) https://BioRender.com/zraz4hk. (d) UMAP depicts ADRB1, ADRB2 and ADRB3 gene expression patterns in total CD8⁺ T cells. Color intensity represents relative gene expression. (e) UMAP plot showing the distribution of naïve and effector/memory CD8⁺ T cells based on their transcriptomic profile, and RNA expression for selected differentiation markers used to distinguish naïve from effector/memory CD8⁺ T cells. (f) ADRB1, ADRB2 and ADRB3 expression level in naïve versus effector/memory CD8⁺ T cells. (g-i) Sub-clustering of effector/memory CD8⁺ T cells into 8 (C0-C7) distinct subsets, and expression level of ADRB2 across C0-C7. Violin colors correspond to cluster identity and dot colors reflect average expression level. (j) Expression of selected markers across C0-C7 used to subcluster effector/memory CD8^+^ T cells into the annotated subsets. (k) Expression of manually selected genes across C0-C7. (l, m) ADRB2 gene expression levels in bystander activated CD8^+^ T cells of patients with acute Hepatitis A Virus (HAV) infection (AHA Bystander, n = 5) and healthy donors (HD Bystander, n = 3) compared to HAV-specific CD8^+^ T cells in the HAV patients (AHA HAV). *p <0.05; **p <0.01; ***p <0.001.

Taken together, ADRB2 is expressed on matched blood and ascites CD8^+^ T cells in patients with decompensated liver cirrhosis. Transcriptomic profiling revealed selective enrichment in bystander compared to antigen-specific CD8^+^ T cells.

### Propranolol attenuates frequency and pro-inflammatory response of bystander CD8⁺ T cells in patients with decompensated liver cirrhosis

To test whether NSBB can directly modulate bystander CD8⁺ T cell function, matched peripheral blood and ascites samples were pre-treated with either the NSBB propranolol or with the ADRB1/ADRB2 agonist isoproterenol for two hours before stimulation with IL-12 + IL-15 + IL-18 or anti-CD3/anti-CD28, respectively, for 24 hours (Fig. 2a). To assess the specific role of ADRB blockade, we here used propranolol, a non-selective ADRB1/ADRB2 antagonist that acts independently of α-adrenergic pathways. Interestingly, cytokine stimulation strongly increased the frequency of bystander-activated CD69⁺CXCR6⁺ CD8⁺ T cells (Fig. 2b, c), whereas pre-treatment with propranolol completely abrogated this effect (Fig. 2b, c). Conversely, co-incubation with isoproterenol resulted in a significant increase in the frequency of bystander-activated CD69^+^CXCR6^+^ CD8^+^ T cells, similar to levels observed in interleukin-stimulated conditions (Fig. 2b, c). Similar to the reduction in CD69^+^CXCR6^+^ CD8^+^ T cell frequencies, pre-treatment with propranolol also downregulated the expression of NKG2D, another marker indicating bystander activation (12), to baseline levels on cytokine-stimulated CD8^+^ T cells in blood and ascites (Fig. 2d).

**Fig. 2.**
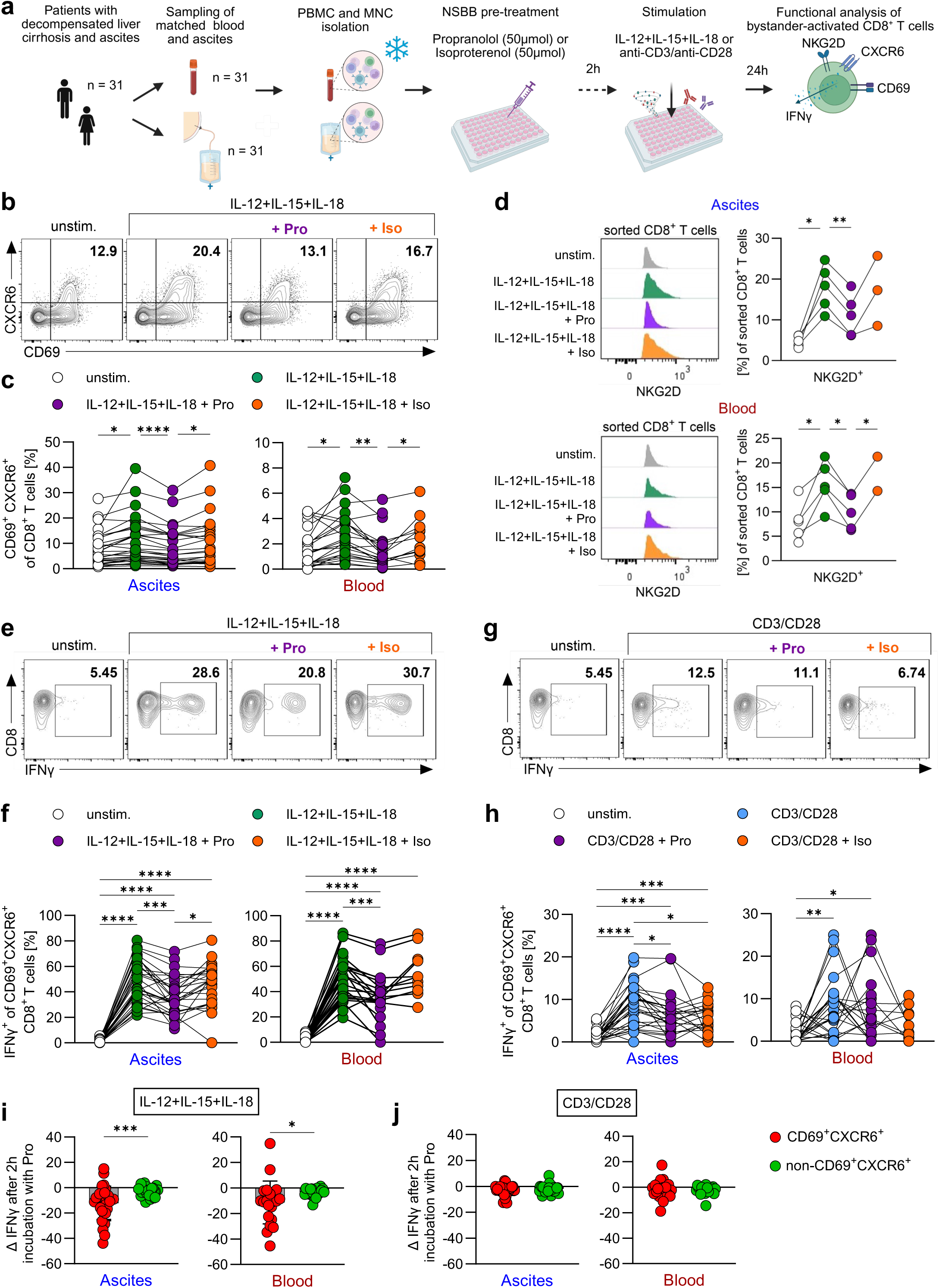
**Propranolol attenuates frequency and pro-inflammatory response of bystander CD8⁺ T cells in patients with decompensated liver cirrhosis.** (a) Schematic workflow of *in vitro* experiments to investigate the impact of NSBB on CD8^+^ T cells. In total, 31 matched ascites and blood samples were obtained. Due to low cell numbers in some samples, 30 ascites samples and 25 blood samples were included in the subsequent analysis. Created in BioRender. Cornberg, M. (2025) https://BioRender.com/8h4p919. (b, c) Representative FACS plots (pre-gated on CD8^+^ T cells) showing the expression of CD69 and CXCR6 on ascites CD8^+^ T cells as well as the respective frequencies of CD69^+^CXCR6^+^ CD8^+^ T cells under unstimulated conditions and after stimulation with IL-12 + IL-15 + IL-18 and pretreatment with propranolol (Pro) or isoproterenol (Iso) in ascites (n = 30) and blood (n = 25) of patients with decompensated cirrhosis. (d) Histogram overlay and quantification of NKG2D expression of sorted CD8^+^ T cells from matched ascites and blood of patients with decompensated cirrhosis (n = 5). (e-h) Representative FACS plots and quantification of IFNγ production in CD69^+^CXCR6^+^ CD8^+^ T cells from ascites (n = 30) and blood (n = 25) of decompensated cirrhosis patients following IL-12 + IL-15 + IL-18 (e, f), or anti-CD3/anti-CD28 (g, h) stimulation ± propranolol or isoproterenol. (i, j) Downregulation of IFNγ production upon propranolol pretreatment in CD69⁺CXCR6⁺ versus non-CD69^+^CXCR6^+^ CD8⁺ T cells from ascites (n = 30) and blood (n = 25) after stimulation with IL-12 + IL-15 + IL-18 (i) or anti-CD3/anti-CD28 (j). Downregulation calculated as Δ (IFNγ⁺ frequency of stimulated cells minus IFNγ⁺ frequency after additional propranolol pretreatment). Wilcoxon signed-rank test for paired comparisons. Unless otherwise specified, multiple comparisons performed with mixed-effects model (REML) with Tukey’s multiple comparisons test. *p <0.05; **p <0.01; ***p <0.001; ****p <0.0001. Schematic figure created with BioRender.com.

To assess the impact of propranolol on the functional profile of bystander-activated (CD69^+^CXCR6^+^) CD8^+^ T cells, we next analyzed the production of pro-inflammatory cytokines and effector molecules following stimulation (Fig. 2e-h, Extended Data Fig. 3c-f). Interestingly, propranolol markedly reduced the frequency of IFNγ^+^ bystander-activated CD8^+^ T cells in both blood and ascites upon cytokine stimulation (Fig. 2e, f). Pre-treatment with isoproterenol was able to restore the IFNγ production in ascites and, to a lesser extent, in blood (Fig. 2e, f). In line with this, TNFα production of bystander-activated CD8^+^ T cells were also diminished by propranolol following cytokine stimulation in ascites and, although not significantly, in blood (Extended Data Fig. 3c, d). In contrast, TCR-mediated (anti-CD3/anti-CD28) IFNγ responses were not affected by propranolol in blood, and only less reduced in ascites (Fig. 2g, h). In addition, no inhibitory effects on CD69^+^CXCR6^+^ CD8^+^ T cells were observed for TNFα (Extended Data Fig. 3e, f). Importantly, propranolol did not impair the cytotoxic potential of blood and ascites CD69^+^CXCR6^+^ CD8^+^ T cells as assessed by Granzyme B, Granzyme K, and CD107a expression after both cytokine-mediated and TCR-mediated stimulation (Extended Data Fig. 3c-f). Noteworthy, CD107a production was even significantly increased following co-incubation with propranolol and cytokine stimulation compared to cytokine stimulation alone in the ascites (Extended Data Fig. 3c). Moreover, the proliferative capacity, indicated by Ki-67 expression, was not impacted by pre-treatment with propranolol and/or isoproterenol (Extended Data Fig. 3c-f).

When investigating the functional responses of the total CD8^+^ T cell compartment, we observed an inhibition of the pro-inflammatory cytokine response (IFNγ and TNFα), whereas the cytotoxic capacity remained unaffected (Granzyme B and Granzyme K) or even increased (CD107a) upon propranolol treatment (Extended Data Fig. 4a-d). Notably, the anti-inflammatory effects of propranolol observed following cytokine stimulation were stronger in bystander-activated (CD69^+^CXCR6^+^) CD8^+^ T cells than in CD8⁺ T cells not co-expressing CD69 and CXCR6 (non-CD69^+^CXCR6^+^) in blood and, particularly, in ascites (Fig. 2i). Following anti-CD3/anti-CD28 stimulation, propranolol had only a minor inhibitory effect that was comparable between the two subsets (Fig. 2j).

Collectively, these results indicate that propranolol reduces the frequency of bystander-activated CD8^+^ T cells and inhibits the pro-inflammatory cytokine response, while only modestly affecting TCR-mediated responses and their cytotoxic potential.

### Preservation of antigen-specific responses upon propranolol treatment is associated with the downregulation of ADRB2 through TCR-mediated signaling

Having established that propranolol inhibits bystander-activated CD8⁺ T cells, we next examined whether it also impairs antigen-specific responses. To this end, we stimulated cells either with innate-like cytokines (IL-12, IL-15, IL-18) or with the CMV pp65 antigen, which induces robust CD8⁺ T cell responses in CMV-seropositive donors (36). As expected, propranolol significantly reduced the IFNγ production of cytokine-stimulated CD8^+^ T cells in blood and ascites (Fig. 3a, b). In contrast, after stimulation with CMV pp65, frequencies of IFNγ^+^ CD8^+^ T cells remained unaffected by propranolol in blood and ascites and were even slightly increased in blood (Fig. 3a, b), indicating preserved antigen-specific CD8⁺ T cell function. Moreover, we performed bulk RNA-sequencing of isolated and differently pre-treated CCR7^-^ CD8^+^ T cells from healthy donors representing effector/memory cells (Fig. 3c), as this subset showed relative upregulation of ADRB2 (Fig. 1e, f). ADRB2 expression was significantly downregulated upon TCR-mediated stimulation with αCD3, aligning with the lower ADRB2 expression level in antigen-specific CD8^+^ T cells as reported above (Fig. 1l, m), whereas stimulation with IL-15 had no significant effect on ADRB2 expression (Fig. 3d). However, αCD3 stimulation together with IL-15 did not reverse ADRB2 downregulation caused by αCD3 (Fig. 3d).

**Fig. 3.**
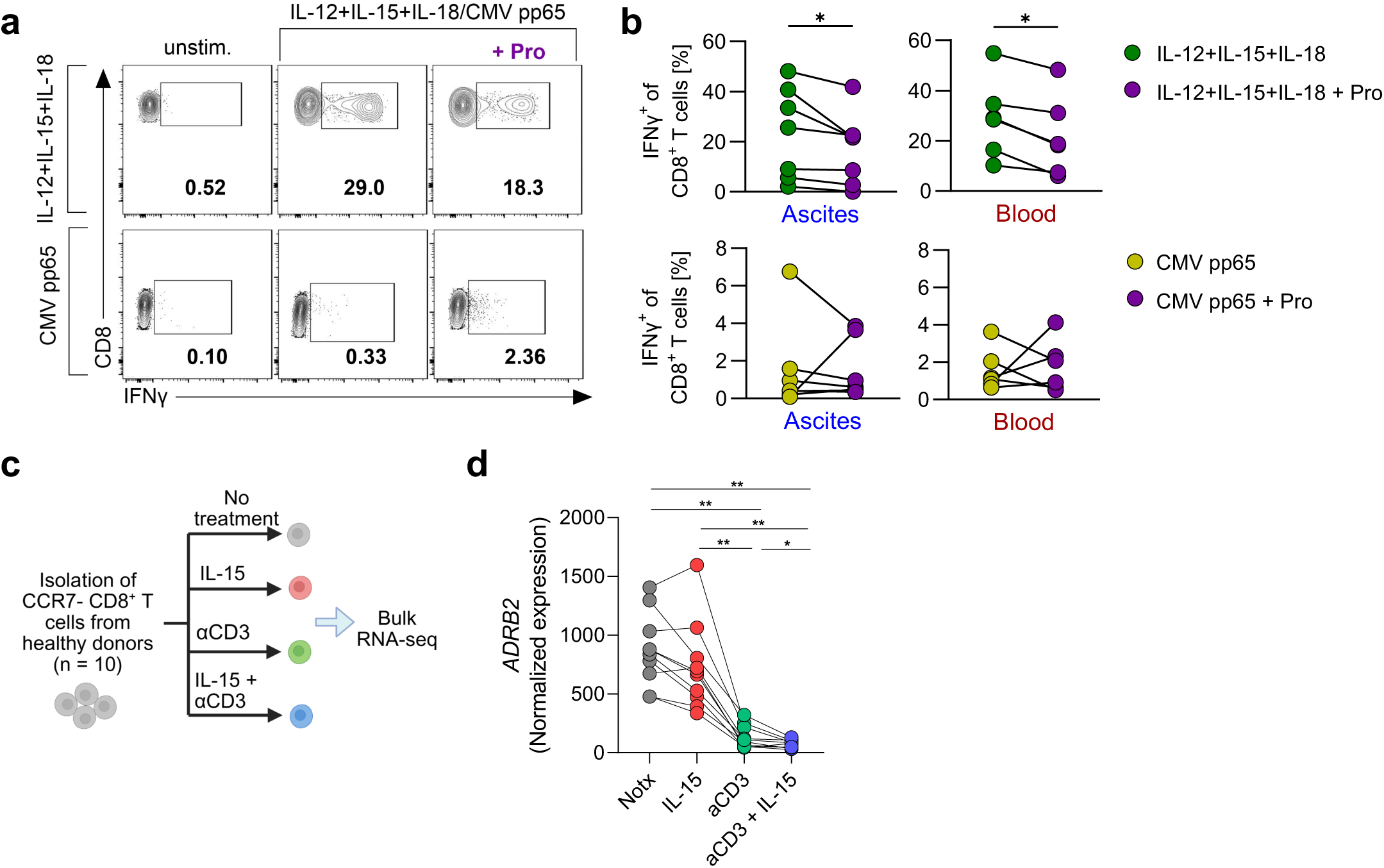
Preservation of antigen-specific responses upon propranolol treatment is associated with the downregulation of ADRB2 through TCR-mediated signaling. (a) Representative FACS plots gated on CD8⁺ T cells in blood, showing IFNγ⁺ cells under unstimulated condition, stimulation with IL-12 + IL-15 + IL-18 or CMV pp65 and pretreatment with propranolol, where indicated. (b) IFNγ production by CD8⁺ T cells from ascites (n = 7) and blood (n = 6) of HLA-A2 positive patients with decompensated cirrhosis following stimulation with IL-12 + IL-15 + IL-18 or CMV pp65, with or without propranolol pretreatment. Wilcoxon signed-rank test. (c) Schematic workflow. Isolated CCR7⁻ CD8⁺ T cells from healthy donors (n = 10) were subjected to four treatment conditions: No treatment (Notx), IL-15, αCD3 and IL-15 + αCD3, followed by bulk RNA-sequencing. Created in BioRender. Cornberg, M. (2025) https://BioRender.com/04t68za. (d) Normalized ADRB2 expression under the indicated experimental conditions. Statistical analysis was performed using Wilcoxon signed-rank test. *p < 0.05; **p < 0.01.

Overall, propranolol reduces the production of pro-inflammatory cytokines by bystander-activated CD8^+^ T cells while maintaining proficient antigen-specific responses. This is consistent with the downregulation of ADRB2 through TCR-mediated signaling.

### Propranolol attenuates bystander-associated transcriptional programs through the STAT1/IRF1 pathway

To investigate the underlying mechanisms by which propranolol reduces pro-inflammatory CD8^+^ T cell responses, we performed bulk RNA-sequencing of sorted CD8^+^ T cells stimulated with IL-12 + IL-15 + IL-18 (IL) or anti-CD3/anti-CD28 (CD), with propranolol (ILP, CDP) or without propranolol (IL, CD) pretreatment (Fig. 4a). Principal component analysis (PCA) revealed that the type of stimulation dominated transcriptional profiles, with propranolol not forming a separate cluster (Fig. 4b). This indicates that propranolol had a smaller impact on the transcriptional features of CD8^+^ T cells than the different stimuli (cytokines or anti-CD3/anti-CD28). However, propranolol induced clear transcriptional changes in both conditions, as evidenced by differently expressed genes (DEGs, Extended Data Fig. 5a). More specifically, pretreatment with propranolol resulted in 699 DEGs (348 up- and 351 downregulated genes) in anti-CD3/anti-CD28 stimulation and 826 DEGs (350 up- and 476 downregulated genes) in cytokine stimulation (Fig. 4c, Extended Data Fig. 5a). The propranolol treatment groups for interleukin and TCR-mediated stimulation shared only 65 DEGs (Fig. 4c), indicating different pharmacological effects depending on the type of stimulation. Furthermore, to investigate potential pathways of propranolol in different CD8^+^ T cell stimulations, we conducted reactome enrichment analysis (Fig. 4d). In general, we observed that propranolol suppressed pathways involved in T cell activation and regulation, whereas no pathways were upregulated upon propranolol treatment. In cytokine-stimulated, but not TCR-stimulated CD8^+^ T cells, propranolol selectively downregulated interferon gamma, alpha/beta, and interferon signaling in general, as well as TNF receptor-ligand binding and the non-canonical NF-kB pathway (Fig. 4d). Consistent with this, transcriptomic analysis revealed that propranolol downregulated signatures associated with interferon signaling and bystander-activation (STAT1, IFNG, interferon (IFN)-regulatory factor (IRF)1, IRF8, IFI44L, MIR155HG, IFIT3, ULBP2) and TNF/TNFR superfamily-mediated signaling (TNF, TNFSF4, TNFSF15, TNFSF18, TNFRSF8, TNFRSF 9; Fig. 4e, f; Extended Data Fig. 5b). Further investigation of the STAT-JAK pathway showed a downregulation of STAT1 and, associated with that, IRF1 after propranolol pre-treatment in cytokine-stimulated, but not TCR-stimulated, cells (Fig. 4f). In parallel, propranolol upregulated NFAM1, previously shown to activate calcineurin-NFAT signaling (37).

**Fig. 4.**
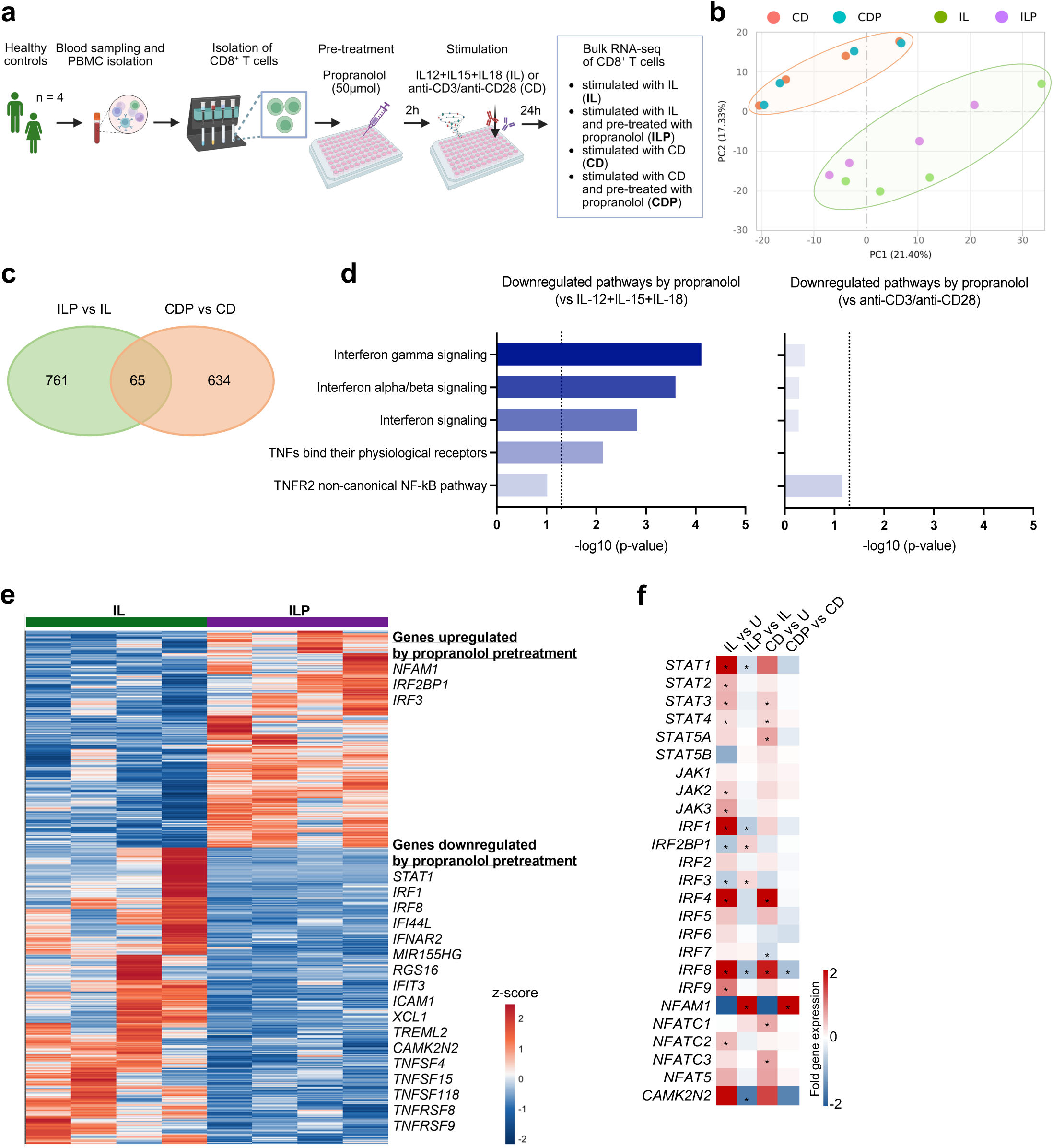
Propranolol attenuates bystander-associated transcriptional programs through the STAT1/IRF1 pathway. (a) Schematic workflow of the experimental setup for bulk RNA-sequencing. Created in BioRender. Cornberg, M. (2025) https://BioRender.com/4puti3z. (b) Principal component analysis (PCA) indicating segregation of the gene expression patterns of sorted CD8^+^ T cells stimulated with IL-12 + IL-15 + IL-18 or anti-CD3/anti-CD28 with or without propranolol pretreatment, respectively, from blood of healthy controls (n = 4). (c) Venn diagram of individual and overlapping differentially expressed genes of CD8^+^ T cells stimulated with IL-12 + IL-15 + IL-18 with/without propranolol pretreatment compared with anti-CD3/anti-CD28 with/without propranolol pretreatment. (d) Analysis of downregulated pathways in DEGs suppressed by propranolol pretreatment following stimulation with IL-12+ IL-15 + IL-18 and anti-CD3/anti-CD28, respectively. Bars indicate −log10 (adjusted p-value) of downregulated pathways. (e) Heatmap of differentially expressed genes between CD8^+^ T cells only stimulated with IL-12 + IL-15 + IL-18 (IL) and CD8^+^ T cells additionally pretreated with propranolol (ILP). (f) Heatmap showing relative up- or downregulation of selected genes associated with JAK-STAT and interferon signaling pathways under cytokine (IL) or anti-CD3/anti-CD28 (CD) stimulation, with or without propranolol pretreatment (ILP, CDP). *p <0.05.

Together, these data identify the STAT1/IRF1 axis as a potential target of propranolol in suppressing cytokine-induced bystander activation.

### NSBB therapy reduces bystander-activated CD8^+^ T cells and ameliorates inflammation in patients with decompensated liver cirrhosis

To validate our *in vitro* observation of reduced bystander CD8^+^ T cell activation following propranolol treatment, we next examined the immunological imprint of NSBB therapy on CD8^+^ T cell phenotypes in patients with decompensated liver cirrhosis. Therefore, the 31 patients included in the *in vitro* analyses were stratified according to their NSBB therapy status: Those receiving NSBB therapy and those who did not (Fig. 5a, Table 1). No significant differences in clinical parameters were observed between NSBB and no-NSBB patients (Table 1). UMAP analysis of paired blood and ascites showed large overlap between CD8^+^ T cells from patients receiving NSBB therapy and patients who did not (Fig. 5b). However, a cluster of CD8^+^ T cells expressing CD69 and CXCR6 could be specifically associated with patients who did not receive NSBB treatment, particularly in blood (Fig. 5b). In line with this, manual gating confirmed significantly lower frequencies of bystander-activated CD8^+^ T cells (CD69⁺CXCR6⁺ and NKG2D⁺) in NSBB-treated patients (Fig. 5c). Notably, this effect was more pronounced in blood than in ascites (Fig. 5c). Other phenotypic markers were comparable between both groups (Extended Data Fig. 6a, b).

**Fig. 5.**
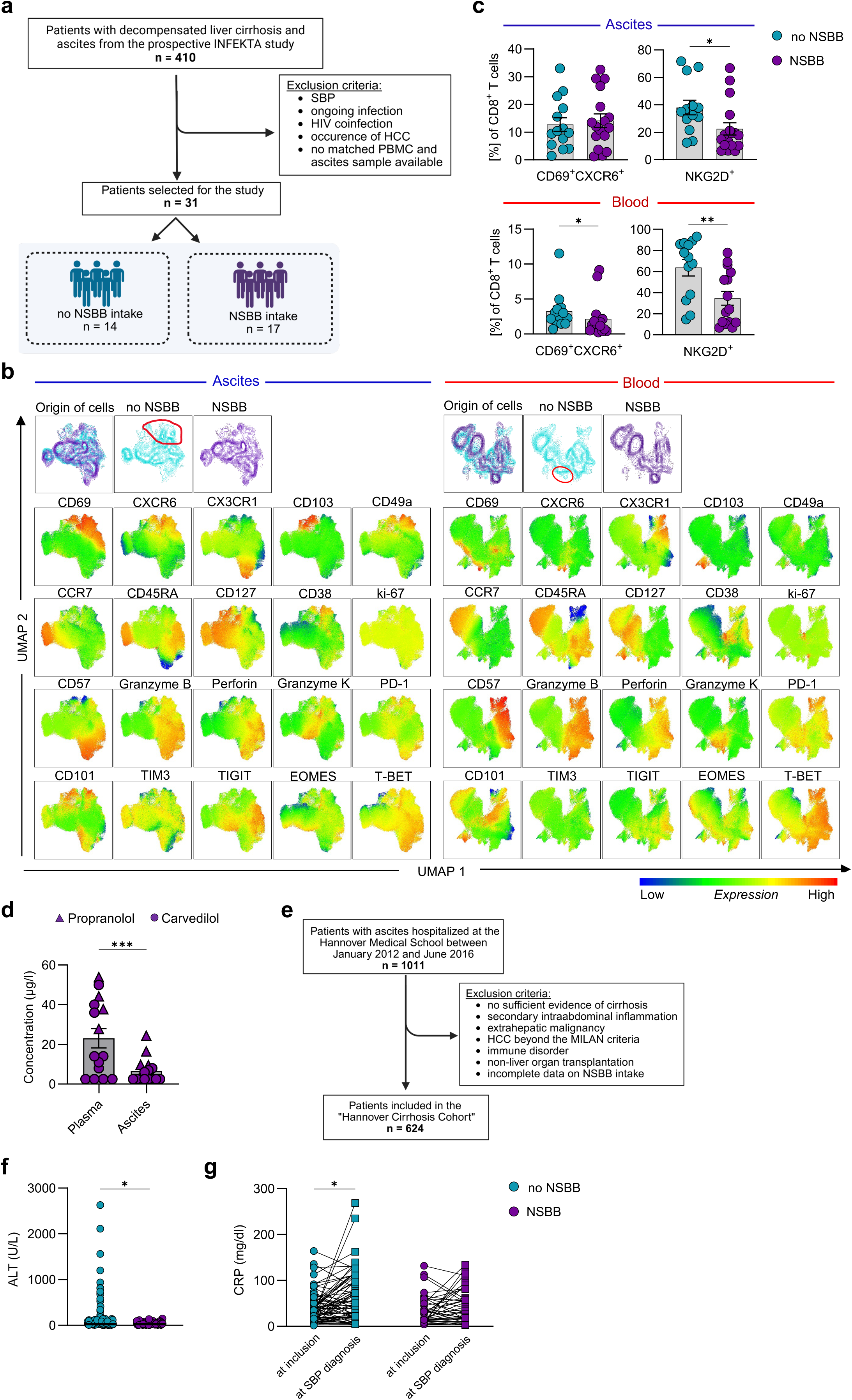
**NSBB therapy reduces bystander-activated CD8^+^ T cells and ameliorates systemic inflammation in patients with decompensated liver cirrhosis.** (a) Flowchart of the experimental study cohort. Patients with decompensated cirrhosis and ascites were recruited from the prospective INFEKTA trial and screened according to predefined exclusion criteria. In total, 31 patients were included and stratified into patients receiving NSBB therapy (n = 17) and patients without NSBB treatment (n = 14). Created in BioRender. Cornberg, M. (2025) https://BioRender.com/rnyyguu. (b) UMAP analysis of CD8^+^ T cells from blood (n = 29) and ascites (n = 31) of patients with and without NSBB treatment. UMAP plots displaying the different treatment groups, and single marker plots depicting expression of indicated phenotypic markers. UMAP analysis was performed on 20 markers for 13 NSBB and non-NSBB patient samples, respectively, with 5000 CD8^+^ T cells exported from each sample. (c) Frequencies of CD69⁺CXCR6⁺ and NKG2D⁺ CD8⁺ T cells in ascites (n = 30-31) and blood (n = 28-29), respectively, determined by *ex vivo* flow cytometry. NKG2D expression was assessed using a separate antibody panel. Unpaired t test (parametric) and Mann-Whitney *U* test (non-parametric). (d) Quantification of propranolol and carvedilol concentrations in plasma and ascites supernatant of NSBB-treated patients (n = 15). Wilcoxon test. (e) Flowchart of the retrospective “Hannover Cirrhosis Cohort” (HAC), previously described by Tergast et al. (38). Created in BioRender. Cornberg, M. (2025) https://BioRender.com/dlo8br8. (f) Alanine aminotransferase (ALT) levels (in U/L) at inclusion (first paracentesis) in patients with NSBB therapy (n = 174) compared to patients without NSBB therapy (n = 304). Mann-Whitney *U* test. (g) C-reactive protein (CRP) values of NSBB (n = 42) compared with no-NSBB patients (n = 74) from the HAC cohort at inclusion versus at the time of diagnosis of spontaneous bacterial peritonitis (SBP). Patients with incomplete laboratory data were excluded from the comparative NSBB analysis. Mann-Whitney *U* test. *p <0.05; **p <0.01; ***p <0.001; ****p <0.0001.

**Table 1.**
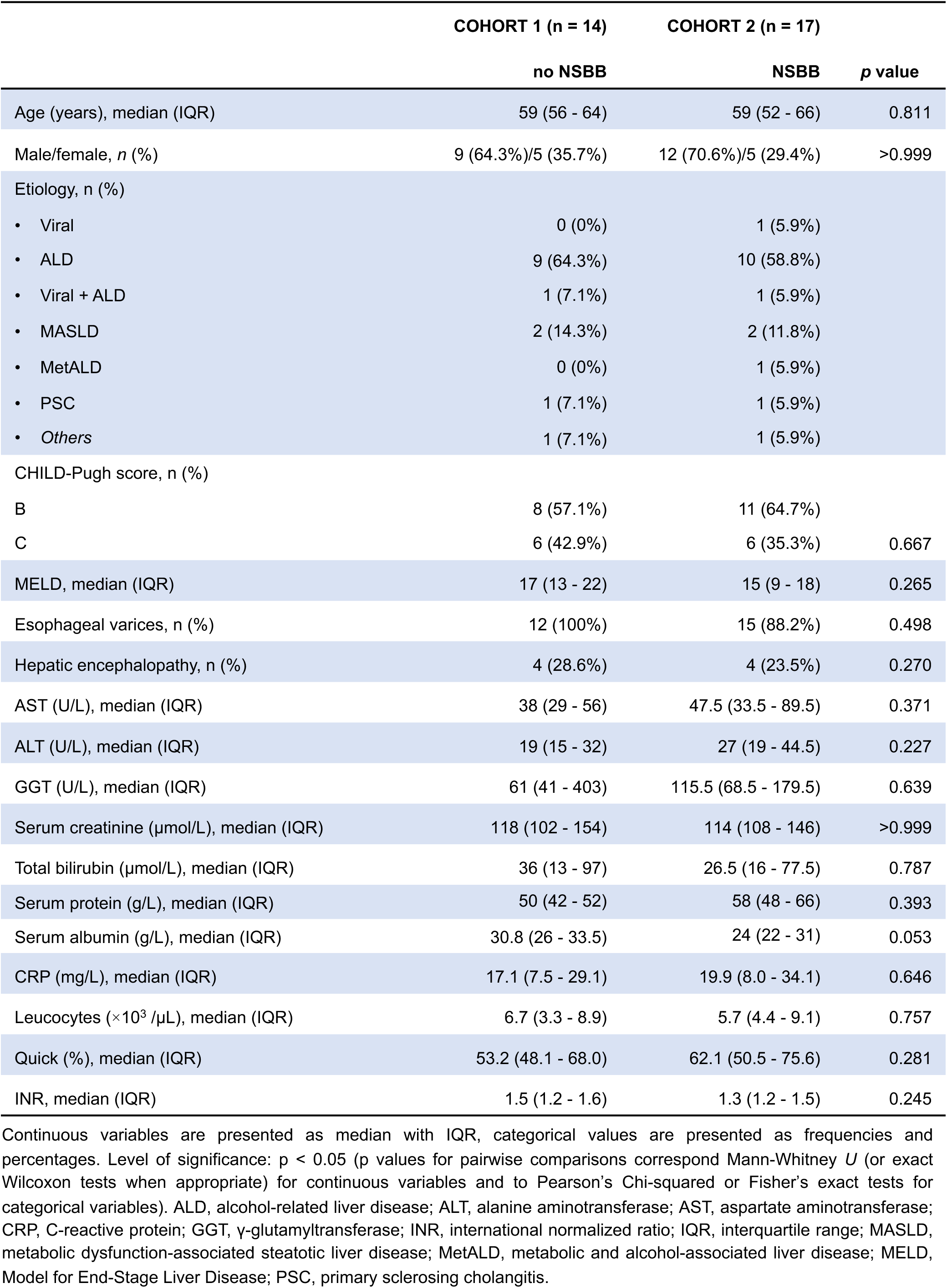
Patients’ clinical characteristics. Continuous variables are presented as median with IQR, categorical values are presented as frequencies and percentages. Level of significance: p < 0.05 (p values for pairwise comparisons correspond to Mann-Whitney *U* (or exact Wilcoxon tests when appropriate) for continuous variables and to Pearson’s Chi-squared or Fisher’s exact tests for categorical variables). ALD, alcohol-related liver disease; ALT, alanine aminotransferase; AST, aspartate aminotransferase; CRP, C-reactive protein; GGT, γ-glutamyltransferase; INR, international normalized ratio; IQR, interquartile range; MASLD, metabolic dysfunction-associated steatotic liver disease; MELD, Model for End-Stage Liver Disease; MetALD, metabolic and alcohol-associated liver disease; PSC, primary sclerosing cholangitis.

|  | COHORT 1 (n = 14) | COHORT 2 (n = 17) |  |
| --- | --- | --- | --- |
|  | no NSBB | NSBB | p value |
| Age (years), median (IQR) | 59 (56 - 64) | 59 (52 - 66) | 0.811 |
| Male/female, n (%) | 9 (64.3%)/5 (35.7%) | 12 (70.6%)/5 (29.4%) | >0.999 |
| Etiology, n (%) |  |  |  |
| • Viral | 0 (0%) | 1 (5.9%) |  |
| • ALD | 9 (64.3%) | 10 (58.8%) |  |
| • Viral + ALD | 1 (7.1%) | 1 (5.9%) |  |
| • MASLD | 2 (14.3%) | 2 (11.8%) |  |
| • MetALD | 0 (0%) | 1 (5.9%) |  |
| • PSC | 1 (7.1%) | 1 (5.9%) |  |
| • Others | 1 (7.1%) | 1 (5.9%) |  |
| CHILD-Pugh score, n (%) |  |  |  |
| B | 8 (57.1%) | 11 (64.7%) |  |
| C | 6 (42.9%) | 6 (35.3%) | 0.667 |
| MELD, median (IQR) | 17 (13 - 22) | 15 (9 - 18) | 0.265 |
| Esophageal varices, n (%) | 12 (100%) | 15 (88.2%) | 0.498 |
| Hepatic encephalopathy, n (%) | 4 (28.6%) | 4 (23.5%) | 0.270 |
| AST (U/L), median (IQR) | 38 (29 - 56) | 47.5 (33.5 - 89.5) | 0.371 |
| ALT (U/L), median (IQR) | 19 (15 - 32) | 27 (19 - 44.5) | 0.227 |
| GGT (U/L), median (IQR) | 61 (41 - 403) | 115.5 (68.5 - 179.5) | 0.639 |
| Serum creatinine (μmol/L), median (IQR) | 118 (102 - 154) | 114 (108 - 146) | >0.999 |
| Total bilirubin (μmol/L), median (IQR) | 36 (13 - 97) | 26.5 (16 - 77.5) | 0.787 |
| Serum protein (g/L), median (IQR) | 50 (42 - 52) | 58 (48 - 66) | 0.393 |
| Serum albumin (g/L), median (IQR) | 30.8 (26 - 33.5) | 24 (22 - 31) | 0.053 |
| CRP (mg/L), median (IQR) | 17.1 (7.5 - 29.1) | 19.9 (8.0 - 34.1) | 0.646 |
| Leucocytes (×10<sup>3</sup> /μL), median (IQR) | 6.7 (3.3 - 8.9) | 5.7 (4.4 - 9.1) | 0.757 |
| Quick (%), median (IQR) | 53.2 (48.1 - 68.0) | 62.1 (50.5 - 75.6) | 0.281 |
| INR, median (IQR) | 1.5 (1.2 - 1.6) | 1.3 (1.2 - 1.5) | 0.245 |

Given the compartment-specific difference, we quantified propranolol and carvedilol concentrations in matched plasma and ascites from treated patients within our NSBB cohort (n = 15). Both drugs were detectable at significant levels, with higher concentrations in blood than in ascites (Fig. 5d). Notably, propranolol but not carvedilol concentrations correlated inversely with frequencies of CD69^+^CXCR6^+^ bystander-activated CD8^+^ T cells (Extended Data Fig. 6c).

To further corroborate our prospective data, we analyzed the large retrospective “Hannover Cirrhosis Cohort”, comprising 624 patients with cirrhosis and ascites, previously published by Tergast et al. (38) (Fig. 5e). Consistent with previous studies (21, 22), NSBB therapy was associated with lower white blood cell (WBC) counts (Extended Data Fig. 6d), as shown earlier (38), as well as lower alanine aminotransferase (ALT) levels (Fig. 5f). Moreover, patients receiving NSBB, who developed spontaneous bacterial peritonitis (SBP), exhibited a significantly blunted rise in C-reactive protein (CRP) during infection compared to no-NSBB patients (Fig. 5g).

Overall, NSBB therapy was associated with reduced frequencies of bystander-activated CD8⁺ T cells. Moreover, patients treated with NSBB exhibited lower levels of inflammation, as indicated by reduced ALT levels, and blunted CRP levels during SBP.

## Discussion

Patients with decompensated cirrhosis and ascites exhibit profound immune dysregulation, a hallmark of cirrhosis-associated immune dysfunction (CAID) (3), with systemic inflammation as a major driver of acute-on-chronic liver failure and mortality (39, 40). Increasing evidence suggests bystander-activated CD8⁺ T cells as key amplifiers of this hyperinflammatory state (8–12). In this study, we identified a direct immunomodulatory effect of NSBB on bystander-activated CD8⁺ T cells and propose a mechanism by which propranolol attenuates systemic inflammation while preserving antigen-specific immunity.

Specifically, we confirmed ADRB2 as the predominant beta-adrenergic receptor on CD8⁺ T cells across different immune compartments (blood, ascites, and liver), with highest expression on effector/memory subsets, in line with previous reports (29–31, 41). We extend these findings by showing that ADRB2 expression is particularly enriched in senescent CD8⁺ T cells, potent producers of pro-inflammatory cytokines (42), and higher in bystander-activated than in antigen-specific CD8⁺ T cells. This implicates a higher sensitivity of bystander CD8^+^ T cells to ADRB2 blockade. Consistent with this, the NSBB propranolol *in vitro* selectively suppressed bystander CD8⁺ T cells, reducing both their frequency (CD69⁺CXCR6⁺, NKG2D⁺) and pro-inflammatory responses following stimulation with IL-12, IL-15 and IL-18, while largely sparing anti-CD3/CD28-mediated responses and preserving antigen-specific (CMV pp65) CD8⁺ T cell immunity. Our findings extend earlier work on bystander CD8⁺ T cell activation in cirrhosis. Previously, we showed that the innate-like cytokines IL-12 and IL-18 potently drive TCR-independent activation of hyperinflammatory CXCR6⁺CD69⁺ CD8⁺ T cells in the ascites (11). *Ibidapo-Obe et al.* recently refined this concept demonstrating that peritoneal macrophages secrete IL-15, which promotes activation of pro-inflammatory CD103⁺CD69⁺ TRM-like CD8⁺ T cells (43). Our present study adds a therapeutic dimension, showing that propranolol interferes with this cytokine-driven circuit through attenuation of harmful bystander responses without impairing antigen-specific immunity. Notably, alongside IFNγ downregulation, we observed increased CD107a expression on CD8^+^ T cells pretreated with propranolol compared with those stimulated by cytokines alone, suggesting a shift towards less inflammatory, but enhanced cytotoxic effector functions. In line with this, a previously published study could show that, in prostate cancer, ADRB2 knockdown in chimeric antigen receptor T cells led to enhanced cytotoxicity against prostate cancer cell lines *in vitro* (44). Mechanistically, propranolol suppressed cytokine-induced activation by downregulating interferon/STAT1/IRF1 signaling, a pathway implicated in cytokine storms and inflammatory pathology through production of IL-12, IL-15, IL-18, IL-23, and nitric oxide (45–47). This is consistent with evidence that ADRB2 signaling through cAMP-PKA can modulate NF-κB and MAPK pathways (48), both of which enhance pro-inflammatory transcription (49, 50) and augment STAT1-dependent transcription (50). While direct evidence for ADRB2-PKA crosstalk with JAK/STAT signaling remains limited, GPCR-JAK/STAT interactions have been reported in other contexts (51). Thus, our data suggest a model in which ADRB2 stimulation amplifies cytokine-induced STAT1 signaling, whereas ADRB2 blockade interrupts this circuit via STAT1, thereby dampening bystander inflammation. In parallel, bulk RNA-seq revealed propranolol-associated induction of NFAM1, a transmembrane adapter protein that enhances calcineurin-NFAT signaling (37). As NFAT activity is regulated post-transcriptionally through dephosphorylation and nuclear translocation rather than transcript abundance (52) and promotes transcriptional programs characteristic of TCR-mediated CD8⁺ T cell activation (53), NFAM1 upregulation under propranolol likely reflects a shift towards a TCR-like activation profile. This is consistent with recent findings that NFATc1 translocation upon TCR engagement suppresses IL-15-driven NKG2D expression and bystander activation in CD8⁺ T cells via AP-1 binding (15). In line with this, we observed that CD8^+^ T cells with elevated expression of ADRB2 exhibit relatively high levels of JUN and FOS (AP-1) (Fig. 1k). Together, this may potentially explain the propranolol-induced downregulation of bystander CD8⁺ T cells observed *in vitro*.

These findings were corroborated *ex vivo* in matched blood and ascites samples showing lower frequencies of bystander-activated (CD69^+^CXCR6^+^ and NKG2D^+^) CD8⁺ T cells in NSBB-treated patients. Notably, the difference between NSBB and no-NSBB patients was more pronounced in blood than in ascites, even though ascites-derived bystander CD8⁺ T cells showed greater sensitivity to propranolol *in vitro*. This may be explained by the lower NSBB concentrations in ascites compared to plasma, which results in a weaker effect on bystander CD8⁺ T cells in the ascites *in vivo.* Consistent with this, propranolol concentrations inversely correlated with bystander CD8⁺ T cell frequencies.

We additionally validated the anti-inflammatory effect of NSBB in a large retrospective cohort. NSBB-treated patients exhibited significantly lower systemic inflammation (WBC) (38) as well as a blunted inflammatory response (CRP) during SBP. These data are consistent with recent studies highlighting the anti-inflammatory effects and improved clinical outcomes associated with NSBB therapy in patients with liver cirrhosis (18–23, 38). However, while these studies mainly attributed the effect to reduced portal hypertension followed by a reduction in intestinal permeability and diminished bacterial translocation (19, 21, 23), our findings suggest an additional, direct effect on immune cells.

More broadly, our study highlights the relevance of the neuro-immune axis in chronic liver disease, where increasing sympathetic neuron signaling exacerbates liver injury and impairs regeneration (54–56). By targeting this axis and selectively restraining bystander CD8⁺ T cell activation via STAT1/IRF1 inhibition, NSBB may attenuate inflammation in cirrhosis, suggesting potential therapeutic benefits even at earlier stages of disease.

In conclusion, this study provides novel insights into a potential mechanism underlying the anti-inflammatory properties of NSBB and supports further exploration of beta-adrenergic blockade as a therapeutic strategy to reduce systemic inflammation while preserving protective immunity in liver cirrhosis.

## Acknowledgements

We thank all study participants, as well as the study nurses and physicians of the Department of Gastroenterology, Hepatology, Infectious Diseases and Endocrinology, Hannover Medical School, for their support in collecting blood and ascites samples. We also thank the laboratory staff and doctoral students for their assistance with biomaterial processing. The manuscript text was refined with the assistance of language editing software.

## Author contributions

Study design: AL, CN, MC. Patient recruitment: MC, AK, BM, TT, HW, ES. Experiments: AL, IT, SK, GZ. Data analysis and statistics: AL, CN, SK, ID, SS, RS, EF. Drafting of the manuscript: AL, CN, MC. Critical revision of the manuscript: all authors.

## Abbreviations

ACLF, acute-on-chronic liver failure; ADRB, beta-adrenergic receptor; ALD, alcohol-related liver disease; ALT, alanine aminotransferase; CAID, cirrhosis-associated immune dysfunction; CMV, cytomegalovirus; DAMPs, damage-associated molecular patterns; DEG, differentially expressed gene; EBV, Epstein-Barr virus; HAC, hepatitis A core protein; HAV, hepatitis A virus; HBV, hepatitis B virus; IAV, influenza A virus; IFN, interferon; IRF, interferon regulatory factor; Iso, isoproterenol-hydrochloride; MASH, metabolic dysfunction-associated steatohepatitis; MASLD, metabolic dysfunction-associated steatotic liver disease; MELD, Model of End-Stage liver disease; MetALD, metabolic and alcohol-associated liver disease; MNC, mononuclear cells; NFAM1, NFAT activating protein with ITAM motif 1; NFAT, nuclear factor of activated T cells; NK cells, natural killer cells; NSBB, non-selective beta-blocker; PAMPs, pathogen-associated molecular patterns; PBMC, peripheral blood mononuclear cells; Pro, propranolol; PSC, primary sclerosing cholangitis; SBP, spontaneous bacterial peritonitis; STAT1, signal transducer and activator of transcription 1; TCR, T cell receptor; TNF, tumor necrosis factor; WBC, white blood cell.

## Methods

### Study design and sample collection

For this study, 36 patients with cirrhosis and ascites decompensation were recruited from the prospective INFEKTA registry (DRKS00010664). The diagnosis of cirrhosis was based on clinical, radiological, or histological findings, and hepatic decompensation was defined by the manifestation of ascites, hepatic encephalopathy, and/or variceal hemorrhage. All patients were hospitalized in the Department of Gastroenterology, Hepatology, Infectious Diseases and Endocrinology at Hannover Medical School. Of the 401 patients with decompensated liver cirrhosis enrolled in the INFEKTA registry, only those with available matched peripheral blood and ascites samples who did not meet any of the exclusion criteria described below were included in this study (n = 36). Samples were collected as part of daily routine, with ascites samples collected when paracentesis was indicated and performed by a physician. Exclusion criteria were manifestation of hepatocellular carcinoma, previous liver transplantation and the presence of SBP, as defined by a cell density of ≥ 250 polymorphonuclear cells/mm^3^ ascites fluid (57). Patients with any other infection at the time of sample acquisition were equally excluded. Furthermore, selected patients (n = 31) were then stratified into patients receiving NSBB therapy (n = 17) and those without NSBB therapy (n = 14). NSBB therapy was initiated by an experienced physician when indicated according to the Baveno criteria (17). Moreover, blood samples of healthy donors (HD, n = 4) were collected for transcriptional analysis.

For analysis of clinical data, the retrospective Hannover Cirrhosis Cohort (HAC), comprising 624 patients with cirrhosis and ascites, was used. The cohort has been described previously in detail (38). For the present study, we performed additional analyses focusing on the impact of NSBB therapy on systemic and hepatic inflammation.

### Isolation and storage of peripheral blood mononuclear cells (PBMC), plasma, mononuclear cells (MNC) from ascites and ascites supernatant

PBMC were isolated from fresh whole blood using standard Ficoll-density gradient separation and erythrocyte lysis buffer. MNC were isolated from ascites by centrifugation at 800 g for 5 min before lysis of red blood cells. Cells were either resuspended in freezing medium (Table 3) and cryopreserved in the gas phase of liquid nitrogen for long-term storage or stained directly after isolation. Plasma was obtained from ethylene diamine tetra acetic acid (EDTA) blood, and ascites supernatant was taken from the ascites sample after centrifugation. Both plasma and ascites supernatant were subsequently stored at -80°C until further use.

### Flow cytometric analysis of PBMC, MNC and isolated CD8^+^ T cells

Cryopreserved PBMC and MNC were thawed and seeded into a 96-well plate at 0.5×10^6^ cells/well. After washing twice in FACS buffer (PBS + 2% FCS + 2 mM EDTA), cells were stained with Live/Dead Fixable Blue or Fixable Viability Stain 700 for 10 minutes in the dark at room temperature to exclude dead cells. Surface and intracellular staining were performed using fluorochrome-labeled monoclonal antibodies (all listed in Extended Data Table S1) for 30 minutes in the dark at 4°C. The FOXP3/Transcription Buffer staining set (eBioscience) or the Cytofix/Cytoperm™ Fixation/Permeabilisation Kit (BD) was used for fixation and permeabilization (Extended Data Table S2). Samples were acquired either on a Cytek Aurora spectral flow cytometer (Cytek Biosciences, USA, CA) or a BD LSRFortessa flow cytometer (BD Biosciences, Germany, Heidelberg). The data obtained were analyzed using FlowJo software V10.10.0. (BD Life Science, USA, NJ). The full gating strategy is displayed in Extended Data Figure 1a. For high-dimensional analysis, Uniform Manifold Approximation and Projection (UMAP) was performed using the open-access FlowJo plugins (58). In selected patients, staining of ADRB1 and ADRB2 was performed on freshly isolated PBMC, as previously described (32).

### Functional assays of PBMC and MNC

For functional analysis of CD8^+^ T cells, either PBMC and MNC or sorted CD8^+^ T cells were stimulated at 0.5×10^6^ cells per well either with IL-12 + IL-15 + IL-18 or purified anti-CD3 and anti-CD28 for 24 hours. Briefly, thawed cells were resuspended in cell culture medium (Extended Data Table S3) and stimulated with IL-12 (10 ng/mL; Miltenyi Biotec, Bergisch Gladbach, Germany) + IL-15 (100 ng/mL; Miltenyi Biotec) + IL-18 (100 ng/mL; MBL International Corporation, Woburn, MA). To stimulate T cell receptor-mediated responses, cells were treated with plate-bound purified anti-CD3 (5µg/mL; Biolegend) and soluble purified anti-CD28 (5µg/mL; Biolegend). For this, 96-well plates were coated with CD3 in PBS overnight at 4°C before cells were seeded and CD28 was added. In selected experiments, CMV pp65 peptide stimulation was performed for 24 hours using 1 µg/mL CMV pp65 (ProImmune, United Kingdom, Oxford), and CMV-specific T cell responses were determined by analyzing the frequencies of IFNγ-producing CD8^+^ T cells. For the last 6 hours of stimulation, Brefeldin A (2 μg/mL; Golgi Plug, BD Biosciences) was added. Simultaneously, staining of CD107a was performed to assess degranulation. In addition, stimulation with PMA and ionomycin was utilized as a positive control.

### *In vitro* treatment with propranolol and isoproterenol

In two distinct experimental setups, either the NSBB propranolol (Pro), an ADRB1 and ADRB2 antagonist (50 µM, P0884, Sigma-Aldrich), or isoproterenol hydrochloride (Iso), an ADRB1 and ADRB2 agonist (50 µM, I6504, Sigma-Aldrich), were added to the cell culture 2 hours prior to stimulation either with IL-12 + IL-15 + IL-18 or anti-CD3/anti-CD28 and incubated at 37°C. The the concentration of propranolol was determined to be effective and non-cytotoxic beforehand (Extended Data. Fig. 3a, b).

### CD8^+^ T cell sorting

In selected experiments, stimulation was performed on sorted blood and ascites CD8^+^ T cells. For this, CD8⁺ T cells were sorted from blood and ascites by fluorescence-activated cell sorting (FACS) (purity confirmed by flow cytometry > 90%, Extended Data. Fig. 1b) or by negative selection using the CD8⁺ T Cell Isolation Kit (Miltenyi Biotec, Bergisch Gladbach, Germany). Non-CD8⁺ cells were labeled with a biotin-conjugated antibody cocktail and subsequently depleted by magnetic separation according to the manufacturer’s instructions.

### Quantification of carvedilol and propranolol concentrations in plasma and ascites supernatant

The concentrations of the NSBB carvedilol and propranolol were measured in matched plasma and ascites supernatant of the patients stratified into the NSBB group (n = 15) by the Bremen Medical Laboratory (Germany). The sample preparation process for carvedilol and propranolol involved protein precipitation using acetonitrile and methanol, followed by dilution in the mobile phase. Analysis was performed using reversed-phase chromatography (Agilent HPLC) in electrospray ionization positive mode on an API4500 triple quadrupole mass spectrometer (AB Sciex) in multiple reaction monitoring (MRM) mode.

### Bulk RNA-sequencing of sorted CD8^+^ T cells

Bulk RNA-sequencing was performed on sorted CD8^+^ T cells from blood of healthy controls (n = 4). After sorting, CD8^+^ T cells were stimulated either with IL-12 + IL-15 + IL-18 or anti-CD3/anti-CD28 for 24 hours with or without propranolol pre-treatment for two hours, and RNA was isolated using the Monarch Spin RNA Isolation Kit #T2110S (New England Biolabs) according to the manufacturer’s instructions. The quantity and integrity of RNA were assessed using a NanoDrop spectrophotometer and a Fragment Analyzer 5400 (Agilent, Santa Clara, CA). Messenger RNA was isolated from total RNA using poly-T-oligo-attached magnetic beads and subsequently fragmented. First- and second-strand cDNA synthesis was then performed.

Following end repair, A-tailing, adapter ligation, size selection, PCR amplification, and purification, the resulting libraries underwent quality control using a Qubit fluorometer (Thermo Fisher Scientific, Waltham, MA), real-time PCR quantification, and a bioanalyzer to assess fragment size distribution. Libraries were pooled and sequenced using Illumina technology.

Raw sequencing data were processed to remove low-quality reads and those containing adapters or poly-N sequences. Reads were aligned to the homo sapiens reference genome (GRCh38/hg38) using HISAT2. Gene expression was quantified using featureCounts (v2.0.6), and expression levels were normalized as FPKM (fragments per kilobase of transcript per million mapped reads). Differential gene expression analysis between treatment groups was conducted using the DESeq2 R package (v1.42.0), applying a threshold of p-value ≤ 0.05.

Differentially expressed genes were visualized using heatmaps generated with the R package pheatmap. Expression values were normalized using row-wise z-score transformation, and hierarchical clustering was performed based on Euclidean distance.

Reactome enrichment analysis was performed using the clusterProfiler R package (v4.8.1). Reactome enrichment pathways with adjusted p-values < 0.05 were considered significantly enriched.

### Analysis of publicly available bulk RNA and single-cell sequencing data

Previously published transcriptomic datasets were reanalyzed in the present study (15). Bulk RNA-sequencing data were derived from isolated CCR7⁻ memory CD8⁺ T cells stimulated with IL-15 (10 ng/ml), anti-CD3 (1 µg/ml), or both for 48 h. Single-cell RNA-sequencing data were obtained from CD8⁺ T cells of healthy donors (n = 3) and patients with acute hepatitis A virus (HAV) infection (n = 5), stained with PE-conjugated MHC-I dCODE dextramers (Immudex) specific for HLA-A*0201 HAV 3D2026 LLYNCCYHV, HLA-A*0201 HCMV pp65495 NLVPMVATV, HLA-A*0201 EBV BMLF1259 GLCTLVAML, and HLA-A*0201 IAV MP58 GILGFVFTL, and sorted into dCODE-positive and -negative populations. Detailed information on library preparation, sequencing, and data processing has been described previously (15).

To compare ADRB1 and ADRB2 expression in circulating and intrahepatic CD8⁺ T cells, we analyzed the single-cell RNA sequencing dataset GSE182159, comprising 46 paired liver and blood samples from 23 individuals across HBV disease states (33). Gene-cell UMI matrices (log-normalized counts per ten thousand) were retrieved from GEO and processed using Trailmaker™, data.table, DropletUtils, and Matrix in R. Low-quality cells (>3% mitochondrial reads, abnormal gene/UMI ratios, or doublets) were excluded. Expression data were normalized (LogNormalize) and batch effects corrected with Harmony. Clustering was performed using the Louvain algorithm (resolution = 0.8) and visualized by UMAP. Differential expression was analyzed with presto (Wilcoxon test, auROC), and group-level comparisons applied a pseudobulk limma-voom workflow.

### Statistical analysis

GraphPad Prism version 10.0.3. (GraphPad Software, San Diego, CA, USA) or R studio (Version 2025.05.1+513) was used for statistical analysis. To assess the distribution of datasets, the D’Agostino & Pearson normality test was used. Paired analysis was performed using paired *t* test for parametric datasets and Wilcoxon signed-rank test for non-parametric datasets. Unpaired datasets were analyzed with unpaired *t* test, if parametric, and Mann-Whitney *U* test, if non-parametric. Comparisons between three or more groups were performed using a mixed-effects model (REML) followed by Tukey’s multiple comparisons test. This approach was used instead of ANOVA due to missing values arising from limited cell numbers, which prevented inclusion of all experimental conditions for every sample. For correlation analysis, either Pearson’s or Spearman’s 𝑟 coefficients were used, depending on the distribution of datasets. Details on the statistical analysis are shown in the respective figure legends. Significant *p*-values are marked as *p <0.05, **p <0.01, ***p <0.001, and ****p <0.0001.

### Study approval

Written informed consent was obtained from all participants before inclusion. The study was conducted in accordance with the ethical guidelines of the 1975 Declaration of Helsinki and was approved by the Ethics Committees of Hannover Medical School (3188-2016; 9227_BO_K_2020, 9474_BO_K_2020, 3193-2016).

## References

1. Devarbhavi H, Asrani SK, Arab JP, Nartey YA, Pose E, Kamath PS. Global burden of liver disease: 2023 update. J Hepatol. 2023;79(2):516–37.

2. Collaborators GC. The global, regional, and national burden of cirrhosis by cause in 195 countries and territories, 1990-2017: a systematic analysis for the Global Burden of Disease Study 2017. Lancet Gastroenterol Hepatol. 2020;5(3):245–66.

3. Albillos A, Martin-Mateos R, Van der Merwe S, Wiest R, Jalan R, Álvarez-Mon M. Cirrhosis-associated immune dysfunction. Nat Rev Gastroenterol Hepatol. 2022;19(2):112–34.

4. Moreau R, Jalan R, Gines P, Pavesi M, Angeli P, Cordoba J, et al. Acute-on-chronic liver failure is a distinct syndrome that develops in patients with acute decompensation of cirrhosis. Gastroenterology. 2013;144(7):1426–37, 37.e1-9.

5. Arroyo V, Moreau R, Jalan R. Acute-on-Chronic Liver Failure. N Engl J Med. 2020;382(22):2137–45.

6. Trebicka J, Fernandez J, Papp M, Caraceni P, Laleman W, Gambino C, et al. PREDICT identifies precipitating events associated with the clinical course of acutely decompensated cirrhosis. J Hepatol. 2021;74(5):1097–108.

7. Costa D, Simbrunner B, Jachs M, Hartl L, Bauer D, Paternostro R, et al. Systemic inflammation increases across distinct stages of advanced chronic liver disease and correlates with decompensation and mortality. J Hepatol. 2021;74(4):819–28.

8. Dudek M, Pfister D, Donakonda S, Filpe P, Schneider A, Laschinger M, et al. Auto-aggressive CXCR6+ CD8 T cells cause liver immune pathology in NASH. Nature. 2021;592(7854):444–9.

9. Nkongolo S, Mahamed D, Kuipery A, Sanchez Vasquez JD, Kim SC, Mehrotra A, et al. Longitudinal liver sampling in patients with chronic hepaSSs B starting antiviral therapy reveals hepatotoxic CD8+ T cells. J Clin Invest. 2023;133(1).

10. Huang CH, Fan JH, Jeng WJ, Chang ST, Yang CK, Teng W, et al. Innate-like bystander-activated CD38+ HLA-DR+ CD8+ T cells play a pathogenic role in patients with chronic hepaSSs C. Hepatology. 2022;76(3):803–18.

11. Niehaus C, Klein S, Strunz B, Freyer E, Maasoumy B, Wedemeyer H, et al. CXCR6+CD69+ CD8+ T cells in the ascites are associated with disease severity in patients with liver cirrhosis. JHEP Reports; 2024.

12. Kefalakes H, Horgan XJ, Jung MK, Amanakis G, Kapuria D, Bolte FJ, et al. Liver-Resident Bystander CD8+ T Cells Contribute to Liver Disease Pathogenesis in Chronic HepaSSs D Virus Infection. Gastroenterology. 2021;161(5):1567–83.e9.

13. Zhang N, Bevan MJ. CD8(+) T cells: foot soldiers of the immune system. Immunity. 2011;35(2):161–8.

14. Balint E, Feng E, Giles EC, Ritchie TM, Qian AS, Vahedi F, et al. Bystander activated CD8+ T cells mediate neuropathology during viral infection via antigen-independent cytotoxicity. Nat Commun. 2024;15(1):896.

15. Lee H, Kim SY, Kim SH, Jeong S, Kim KH, Kim CG, et al. TCR signaling via NFATc1 constrains IL-15-induced bystander activation of human memory CD8+ T cells. Immunity. 2025.

16. Liver. EAitiot. EASL Clinical Practice Guidelines for the management of patients with decompensated cirrhosis. J Hepatol. 2018;69(2):406–60.

17. de Franchis R, Bosch J, Garcia-Tsao G, Reiberger T, Ripoll C, Faculty BV. Baveno VII - Renewing consensus in portal hypertension. J Hepatol. 2022;76(4):959–74.

18. Villanueva C, Albillos A, Genescà J, Garcia-Pagan JC, Calleja JL, Aracil C, et al. β blockers to prevent decompensation of cirrhosis in patients with clinically significant portal hypertension (PREDESCI): a randomised, double-blind, placebo-controlled, multicentre trial. Lancet. 2019;393(10181):1597–608.

19. Reiberger T, Ferlitsch A, Payer BA, Mandorfer M, Heinisch BB, Hayden H, et al. Non-selective betablocker therapy decreases intestinal permeability and serum levels of LBP and IL-6 in patients with cirrhosis. J Hepatol. 2013;58(5):911–21.

20. Mookerjee RP, Pavesi M, Thomsen KL, Mehta G, Macnaughtan J, Bendtsen F, et al. Treatment with non-selective beta blockers is associated with reduced severity of systemic inflammation and improved survival of patients with acute-on-chronic liver failure. J Hepatol. 2016;64(3):574–82.

21. Jachs M, Hartl L, Schaufler D, Desbalmes C, Simbrunner B, Eigenbauer E, et al. Amelioration of systemic inflammation in advanced chronic liver disease upon beta-blocker therapy translates into improved clinical outcomes. Gut. 2021;70(9):1758–67.

22. Kumar R, Lin S, Mehta G, Mesquita MD, Calvao JAF, Sheikh MF, et al. Non-selective beta-blocker is associated with reduced mortality in critically ill patients with cirrhosis: A real-world study. Aliment Pharmacol Ther. 2024.

23. Albillos A, Krag A. Beta-blockers in the era of precision medicine in patients with cirrhosis. J Hepatol. 2023;78(4):866–72.

24. Sanders VM. The beta2-adrenergic receptor on T and B lymphocytes: do we understand it yet? Brain Behav Immun. 2012;26(2):195–200.

25. Sanders VM, Baker RA, Ramer-Quinn DS, Kasprowicz DJ, Fuchs BA, Street NE. Differential expression of the beta2-adrenergic receptor by Th1 and Th2 clones: implications for cytokine production and B cell help. J Immunol. 1997;158(9):4200–10.

26. Tsai HC, Hsu CF, Huang CC, Huang SF, Li TH, Yang YY, et al. Propranolol Suppresses the T-Helper Cell Depletion-Related Immune Dysfunction in Cirrhotic Mice. Cells. 2020;9(3).

27. Takenaka MC, Araujo LP, Maricato JT, Nascimento VM, Guereschi MG, Rezende RM, et al. Norepinephrine Controls Effector T Cell DifferentiaSon through β2-Adrenergic Receptor-Mediated Inhibition of NF-κB and AP-1 in Dendritic Cells. J Immunol. 2016;196(2):637–44.

28. Jürgens M, Claus M, Wingert S, Niemann JA, Picard LK, Hennes E, et al. Chronic stimulation desensitizes β2-adrenergic receptor responses in natural killer cells. Eur J Immunol. 2024:e2451299.

29. Slota C, Shi A, Chen G, Bevans M, Weng NP. Norepinephrine preferentially modulates memory CD8 T cell function inducing inflammatory cytokine production and reducing proliferation in response to activation. Brain Behav Immun. 2015;46:168–79.

30. Estrada LD, Ağaç D, Farrar JD. Sympathetic neural signaling via the β2-adrenergic receptor suppresses T-cell receptor-mediated human and mouse CD8(+) T-cell effector function. Eur J Immunol. 2016;46(8):1948–58.

31. Daher C, Vimeux L, Stoeva R, Peranzoni E, Bismuth G, Wieduwild E, et al. Blockade of β-Adrenergic Receptors Improves CD8^+^ T-cell Priming and Cancer Vaccine Efficacy. Cancer Immunol Res. 2019;7(11):1849–63.

32. Globig AM, Zhao S, Roginsky J, Maltez VI, Guiza J, Avina-Ochoa N, et al. The β1-adrenergic receptor links sympathetic nerves to T cell exhaustion. Nature. 2023;622(7982):383–92.

33. Zhang C, Li J, Cheng Y, Meng F, Song JW, Fan X, et al. Single-cell RNA sequencing reveals intrahepatic and peripheral immune characteristics related to disease phases in HBV-infected patients. Gut. 2023;72(1):153–67.

34. Kim J, Chang DY, Lee HW, Lee H, Kim JH, Sung PS, et al. Innate-like Cytotoxic Function of Bystander-Activated CD8^+^ T Cells Is Associated with Liver Injury in Acute HepaSSs A. Immunity. 2018;48(1):161–73.e5.

35. Schulte I, Hitziger T, Giugliano S, Timm J, Gold H, Heinemann FM, et al. Characterization of CD8^+^ T-cell response in acute and resolved hepaSSs A virus infection. J Hepatol. 2011;54(2):201–8.

36. Kern F, Bunde T, Faulhaber N, Kiecker F, Khatamzas E, Rudawski IM, et al. Cytomegalovirus (CMV) phosphoprotein 65 makes a large contribution to shaping the T cell repertoire in CMV-exposed individuals. J Infect Dis. 2002;185(12):1709–16.

37. Yang J, Hu G, Wang SW, Li Y, Martin R, Li K, et al. Calcineurin/nuclear factors of activated T cells (NFAT)-activating and immunoreceptor tyrosine-based activation motif (ITAM)-containing protein (CNAIP), a novel ITAM-containing protein that activates the calcineurin/NFAT-signaling pathway. J Biol Chem. 2003;278(19):16797–801.

38. Tergast TL, Kimmann M, Laser H, Gerbel S, Manns MP, Cornberg M, et al. Systemic arterial blood pressure determines the therapeutic window of non-selective beta blockers in decompensated cirrhosis. Aliment Pharmacol Ther. 2019;50(6):696–706.

39. Arroyo V, Angeli P, Moreau R, Jalan R, Clària J, Trebicka J, et al. The systemic inflammation hypothesis: Towards a new paradigm of acute decompensation and multiorgan failure in cirrhosis. J Hepatol. 2021;74(3):670–85.

40. Clària J, Stauber RE, Coenraad MJ, Moreau R, Jalan R, Pavesi M, et al. Systemic inflammation in decompensated cirrhosis: Characterization and role in acute-on-chronic liver failure. Hepatology. 2016;64(4):1249–64.

41. Qiao G, Chen M, Mohammadpour H, MacDonald CR, Bucsek MJ, Hylander BL, et al. Chronic Adrenergic Stress Contributes to Metabolic Dysfunction and an Exhausted Phenotype in T Cells in the Tumor Microenvironment. Cancer Immunol Res. 2021;9(6):651–64.

42. Zhang J, He T, Xue L, Guo H. Senescent T cells: a potential biomarker and target for cancer therapy. EBioMedicine. 2021;68:103409.

43. Ibidapo-Obe O, Stengel S, Frissen M, Reißing J, Große K, Rooney M, et al. Macrophage-derived IL-15 imprints peritoneal TRM-like CD8 T cells in cirrhosis and spontaneous bacterial peritonitis. JHEP Rep. 2025;7(6):101381.

44. Ajmal I, Farooq MA, Duan Y, Yao J, Gao Y, Hui X, et al. Intrinsic ADRB2 inhibition improves CAR-T cell therapy efficacy against prostate cancer. Mol Ther. 2024;32(10):3539–57.

45. Sundaram B, Pandian N, Mall R, Wang Y, Sarkar R, Kim HJ, et al. NLRP12-PANoptosome activates PANoptosis and pathology in response to heme and PAMPs. Cell. 2023;186(13):2783–801.e20.

46. Lohoff M, Mak TW. Roles of interferon-regulatory factors in T-helper-cell differentiaSon. Nat Rev Immunol. 2005;5(2):125–35.

47. Wang L, Zhu Y, Zhang N, Xian Y, Tang Y, Ye J, et al. The multiple roles of interferon regulatory factor family in health and disease. Signal Transduct Target Ther. 2024;9(1):282.

48. Lorton D, Bellinger DL. Molecular mechanisms underlying β-adrenergic receptor-mediated cross-talk between sympathetic neurons and immune cells. Int J Mol Sci. 2015;16(3):5635–65.

49. Christian F, Smith EL, Carmody RJ. The Regulation of NF-κB Subunits by Phosphorylation. Cells. 2016;5(1).

50. Ramsauer K, Sadzak I, Porras A, Pilz A, Nebreda AR, Decker T, et al. p38 MAPK enhances STAT1-dependent transcription independently of Ser-727 phosphorylation. Proc Natl Acad Sci U S A. 2002;99(20):12859–64.

51. Sasaguri T, Teruya H, Ishida A, Abumiya T, Ogata J. Linkage between alpha(1) adrenergic receptor and the Jak/STAT signaling pathway in vascular smooth muscle cells. Biochem Biophys Res Commun. 2000;268(1):25–30.

52. Okamura H, Aramburu J, García-Rodríguez C, Viola JP, Raghavan A, Tahiliani M, et al. Concerted dephosphorylation of the transcription factor NFAT1 induces a conformational switch that regulates transcriptional activity. Mol Cell. 2000;6(3):539–50.

53. Macian F. NFAT proteins: key regulators of T-cell development and function. Nat Rev Immunol. 2005;5(6):472–84.

54. Zhou Y, Lin X, Jiao Y, Yang D, Li Z, Zhu L, et al. A brain-to-liver signal mediates the inhibition of liver regeneration under chronic stress in mice. Nat Commun. 2024;15(1):10361.

55. Sharma S, Le Guillou D, Chen JY. Cellular stress in the pathogenesis of nonalcoholic steatohepaSSs and liver fibrosis. Nat Rev Gastroenterol Hepatol. 2023;20(10):662–78.

56. Xu MY, Guo CC, Li MY, Lou YH, Chen ZR, Liu BW, et al. Brain-gut-liver axis: Chronic psychological stress promotes liver injury and fibrosis. Front Cell Infect Microbiol. 2022;12:1040749.

57. Gerbes AL, Gülberg V, Sauerbruch T, Wiest R, Appenrodt B, Bahr MJ, et al. [German S 3-guideline "ascites, spontaneous bacterial peritonitis, hepatorenal syndrome"]. Z Gastroenterol. 2011;49(6):749–79.

58. Becht E, McInnes L, Healy J, Dutertre CA, Kwok IWH, Ng LG, et al. Dimensionality reduction for visualizing single-cell data using UMAP. Nat Biotechnol. 2018.

