## Extended Data for "Non-selective beta-blockers reduce bystander CD8^+^ T cell activation in decompensated liver cirrhosis"

**Table of Contents**

Extended Data Figure Legends

Extended Data Figure 1

Extended Data Figure 2

Extended Data Figure 3

Extended Data Figure 4

Extended Data Figure 5

Extended Data Figure 6

Extended Data Table S1

Extended Data Table S2

Extended Data Table S3

**Extended Data Figure 1. Gating strategy.**

(a) Gating strategy for identification of bystander-activated CD69$^+$CXCR6$^+$ CD8$^+$ T cells. (b) Representative flow cytometry plots showing sorting quality of CD8$^+$ T cells, indicated by CD8$^+$ T cell gating before (upper row) and after (lower row) cell sorting.

**Extended Data Figure 2. Expression of ADRB1 and ADRB2 on CD8$^+$ T cells in blood, ascites, and liver.**

(a) Expression of ADRB1 and ADRB2 (mean fluorescence intensity, MFI) on CD8$^+$ T cells from paired blood and ascites samples (n = 4). Wilcoxon signed-rank test. (b) UMAP visualization of CD8$^+$ T cells derived from previously published single-cell RNA-sequencing data (GSE182159) from paired liver and blood samples of individuals with different HBV infection states. (c, d) UMAP visualizations showing expression levels of ADRB1 (c) and ADRB2 (d) in CD8$^+$ T cells across liver and blood. (e, f) Violin plots displaying expression of ADRB1 (e) and ADRB2 (f) in paired blood and liver-derived CD8$^+$ T cells.

**Extended Data Figure 3. Functional analysis of bystander-activated (CD69$^+$CXCR6$^+$) CD8$^+$ T cells.**

(a, b) Frequencies of living CD8$^+$ T cells upon pretreatment with propranolol compared to IL-12 + IL-15 + IL-18 stimulated condition in (a) ascites (n = 29) and (b) blood (n = 22). Wilcoxon signed-rank test. (c, d) Functional responses of CD69$^+$CXCR6$^+$ CD8$^+$ T cells following cytokine stimulation, with propranolol or isoproterenol pretreatment where indicated, in (c) ascites (n = 29-30) and (d) blood (n = 24-26). (e, f) Functional responses of CD69$^+$CXCR6$^+$ CD8$^+$ T cells after anti-CD3/anti-CD28 stimulation, with propranolol or isoproterenol pretreatment where indicated, in (e) ascites (n = 26-29) and (f) blood (n = 12-15). Mixed-effects model (REML) with Tukey's multiple comparisons test. *p <0.05; **p <0.01; ***p <0.001; ****p <0.0001.

**Extended Data Figure 4. Functional analysis of total $CD8^+$ T cells.**

(a, b) Functional readout of total $CD8^+$ T cells following cytokine stimulation, with propranolol or isoproterenol pretreatment where indicated, in (a) ascites (n = 29) and (b) blood (n = 22). (c, d) Functional readout of total $CD8^+$ T cells stimulated with anti-CD3/anti-CD28 with propranolol or isoproterenol pretreatment where indicated in (c) ascites (n = 26-29) and (d) blood (n = 12-15). Mixed-effects model (REML) with Tukey's multiple comparisons test. *p <0.05; **p <0.01; ***p <0.001; ****p <0.0001.

**Extended Data Figure 5. Bulk sequencing of $CD8^+$ T cells after propranolol treatment.**

(a) Counts of differently expressed genes (ALL), upregulated (UP) and downregulated (DOWN) genes. (b) Heatmap showing the z-score (red = positive z-score; blue = negative z-score) of differentially expressed genes between $CD8^+$ T cells only stimulated with anti-CD3/anti-CD28 (CD) and $CD8^+$ T cells additionally pretreated with propranolol (CDP).

**Extended Data Figure 6. Phenotypic analysis of $CD8^+$ T cells in NSBB-treated compared to untreated patients and correlation with NSBB levels.**

(a, b) Comparison of phenotypic marker expression on total $CD8^+$ T cells between NSBB patients (n = 17) and patients not receiving NSBB (n = 13) in (a) ascites and (b) blood. Multiple Mann-Whitney tests. (c) Correlation of blood and ascites propranolol (n = 4) and carvedilol (n = 11) concentrations with *ex vivo* measured $CD69^+CXCR6^+$ $CD8^+$ T cell frequencies. Spearman's $r$ coefficient. (d) White blood cell (WBC) counts (in 1000/µl) at inclusion (first paracentesis) in patients with NSBB therapy (n = 194) compared to patients without NSBB therapy (n = 318). Patients with incomplete laboratory data were excluded from the comparative NSBB analysis. Mann-Whitney *U* test. *p <0.05; **p <0.01; ***p <0.001; ****p <0.0001.

Extended Data Figure 1. Gating strategy.

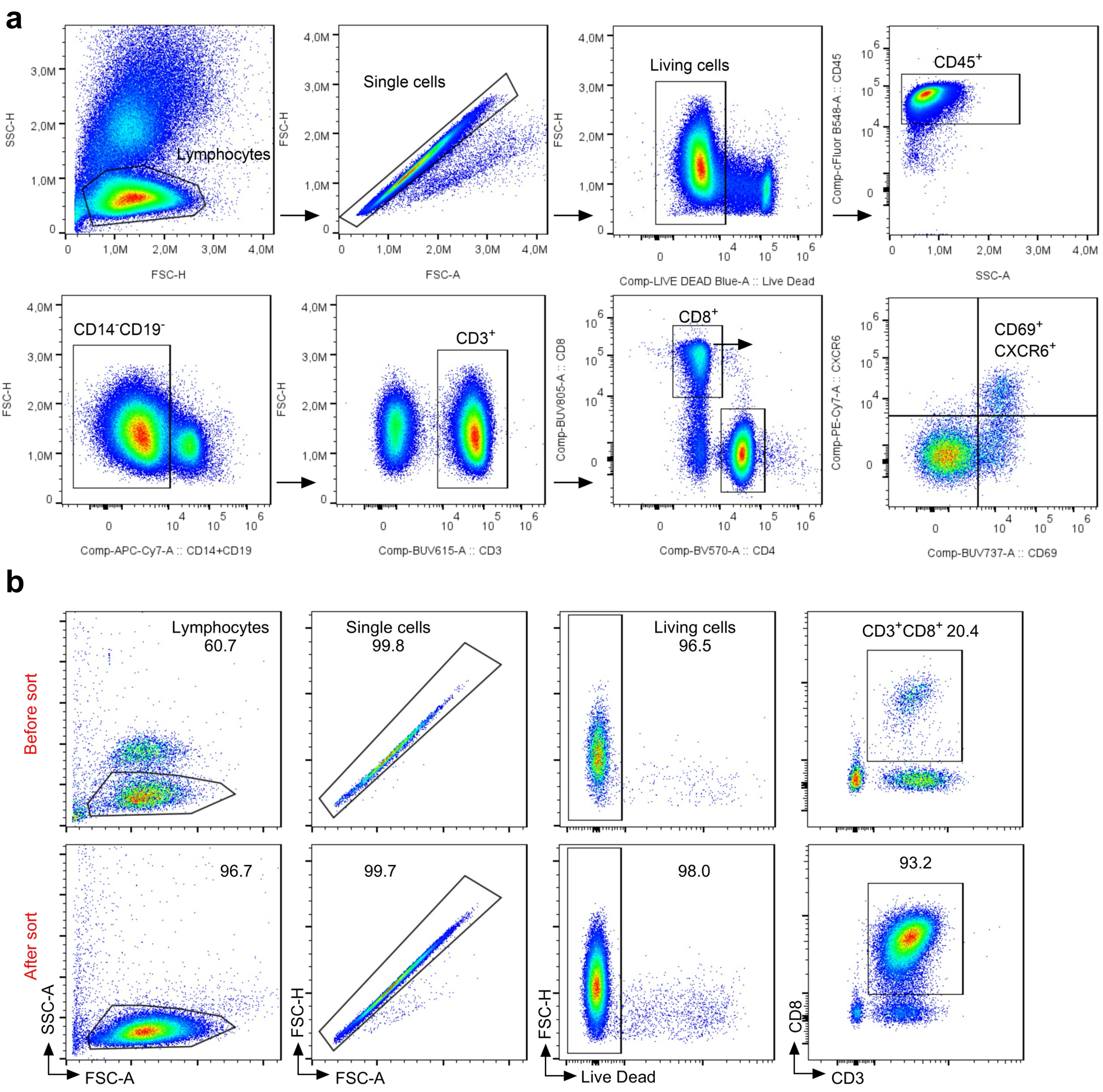

Extended Data Figure 2. Expression of ADRB1 and ADRB2 on CD8⁺ T cells in blood, ascites, and liver.

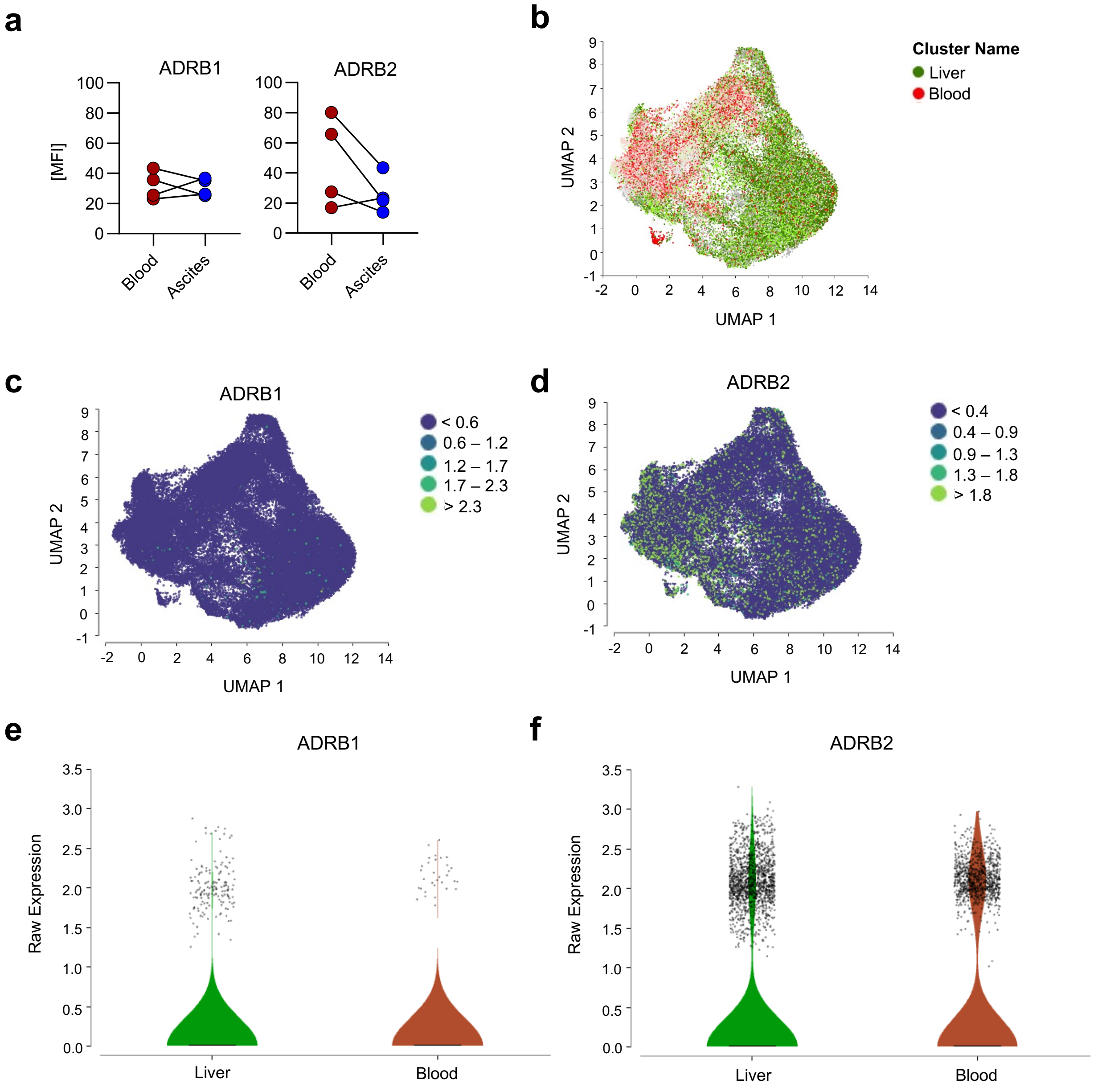

Extended Data Figure 3. Functional analysis of bystander-activated (CD69⁺CXCR6⁺) CD8⁺ T cells.

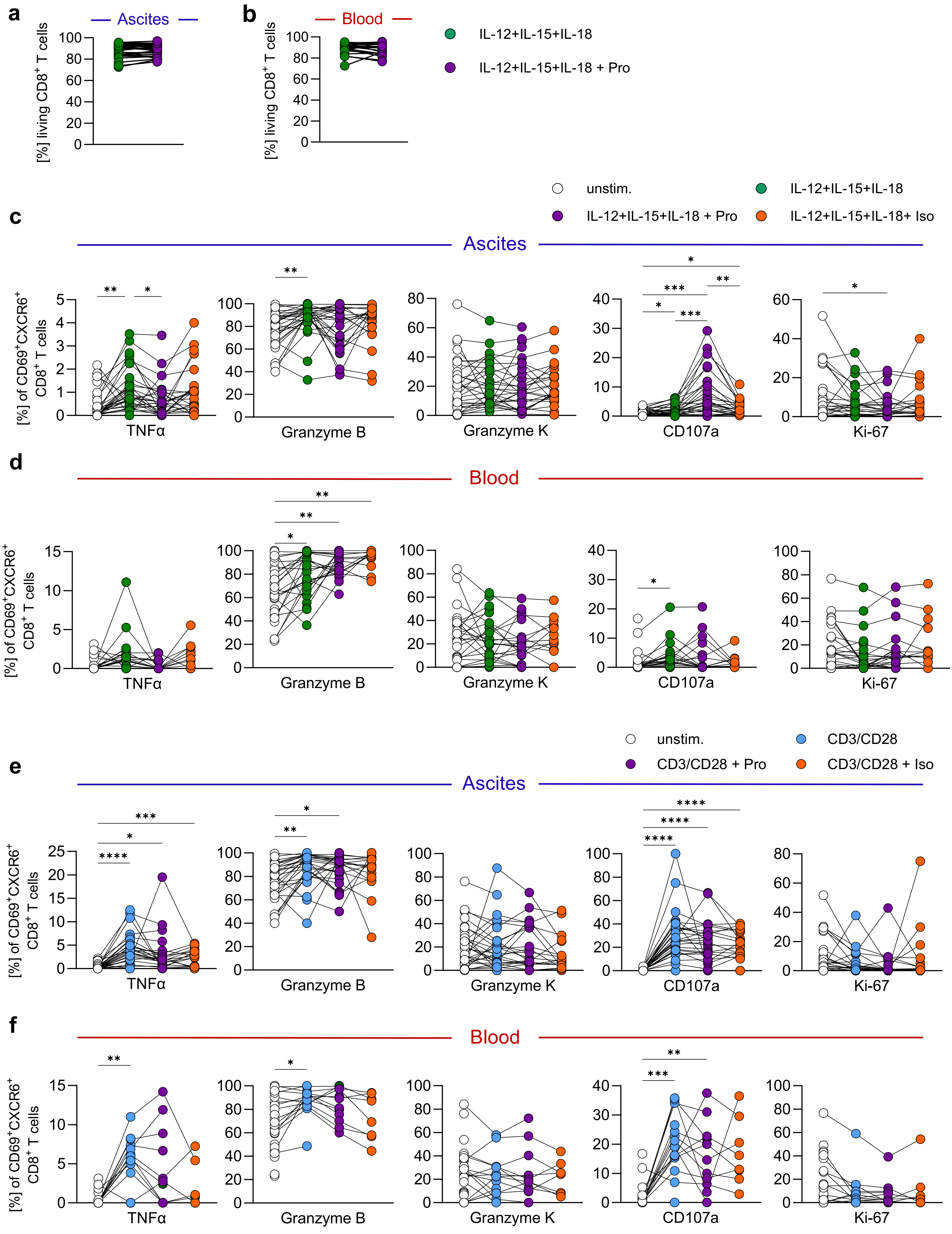

Extended Data Figure 4. Functional analysis of total CD8⁺ T cells.

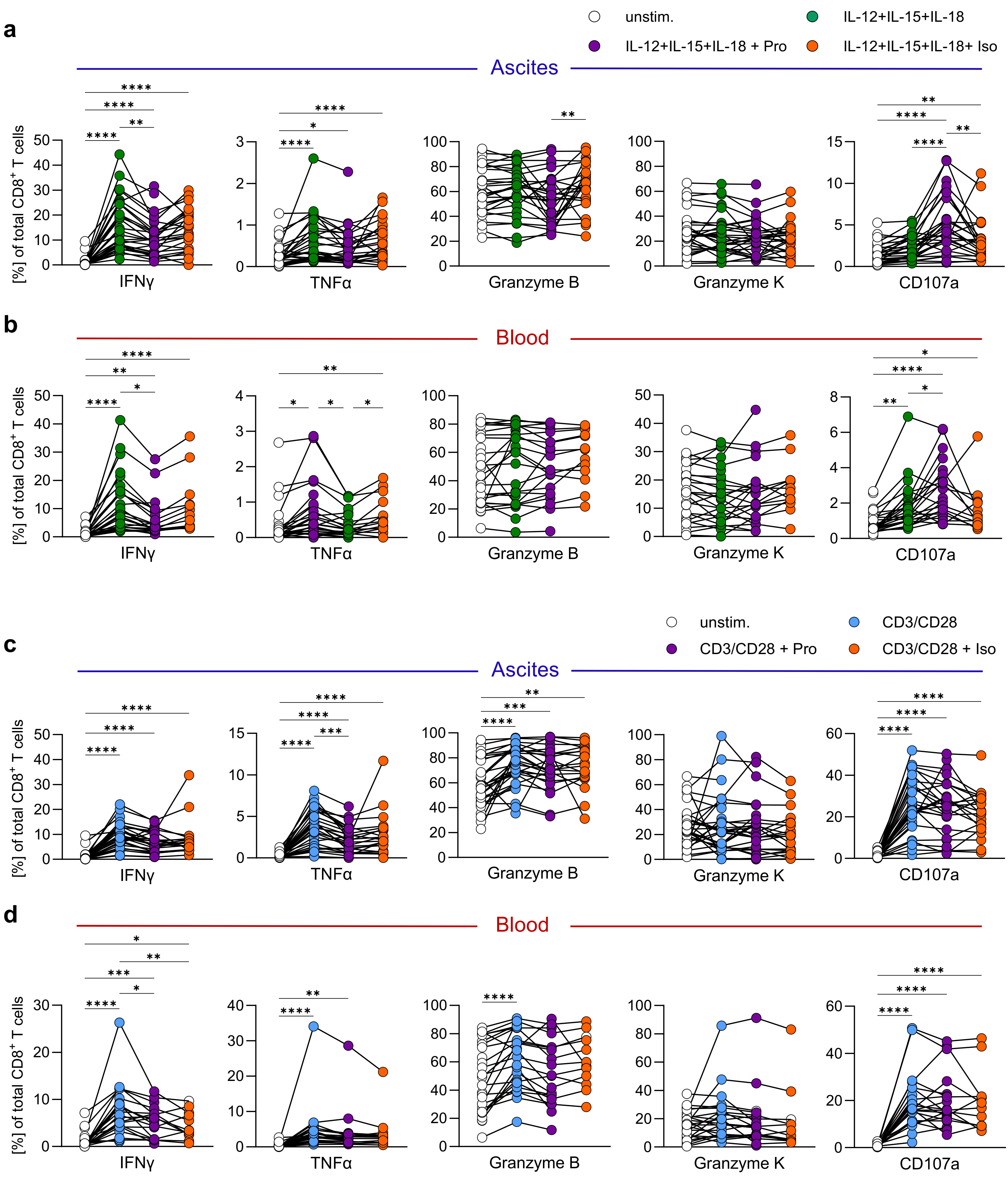

Extended Data Figure 5. Bulk sequencing of CD8⁺ T cells after propranolol treatment.

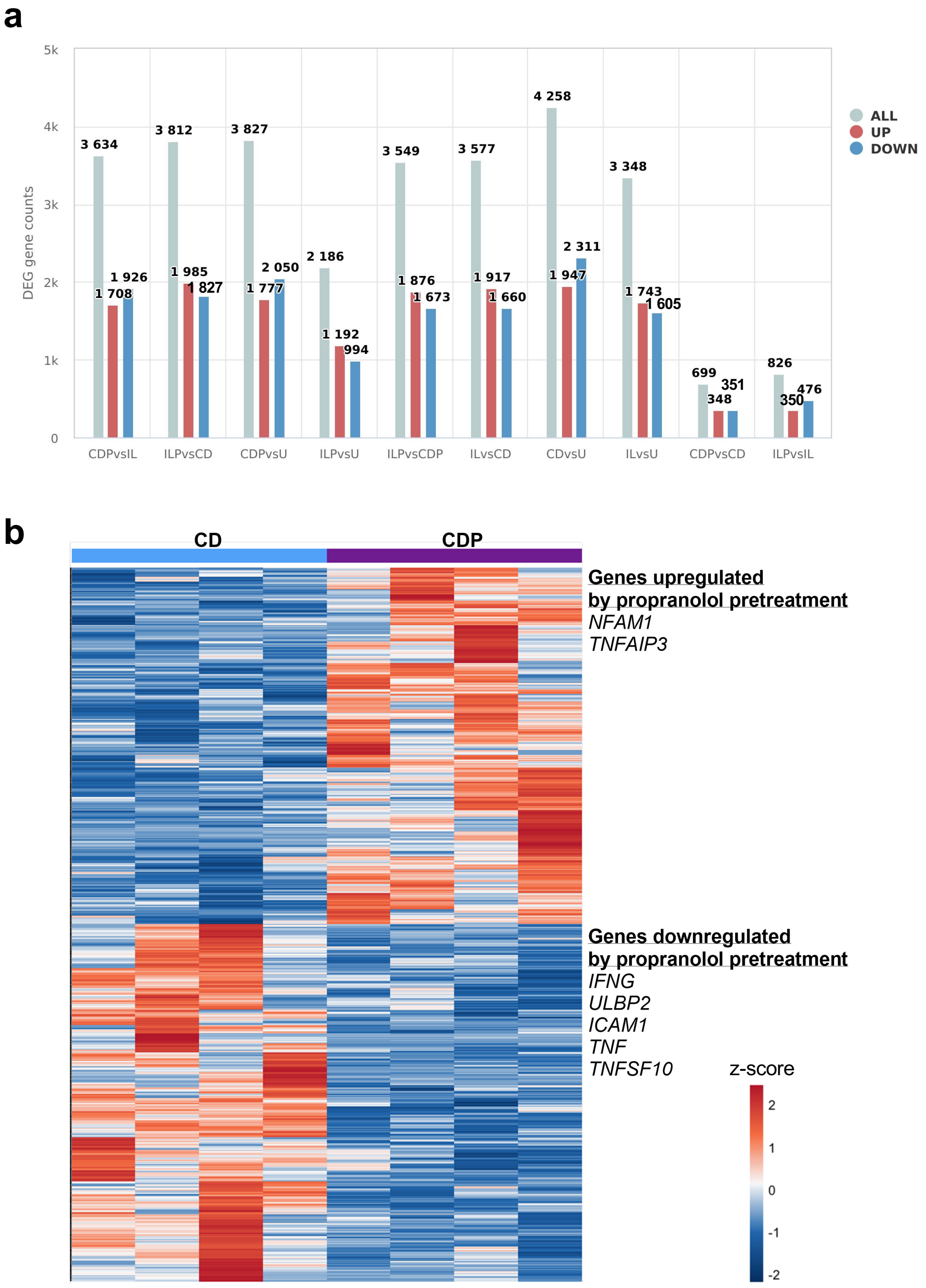

Extended Data Figure 6. Phenotypic analysis of CD8⁺ T cells in NSBB-treated compared to untreated patients and correlation with NSBB levels.

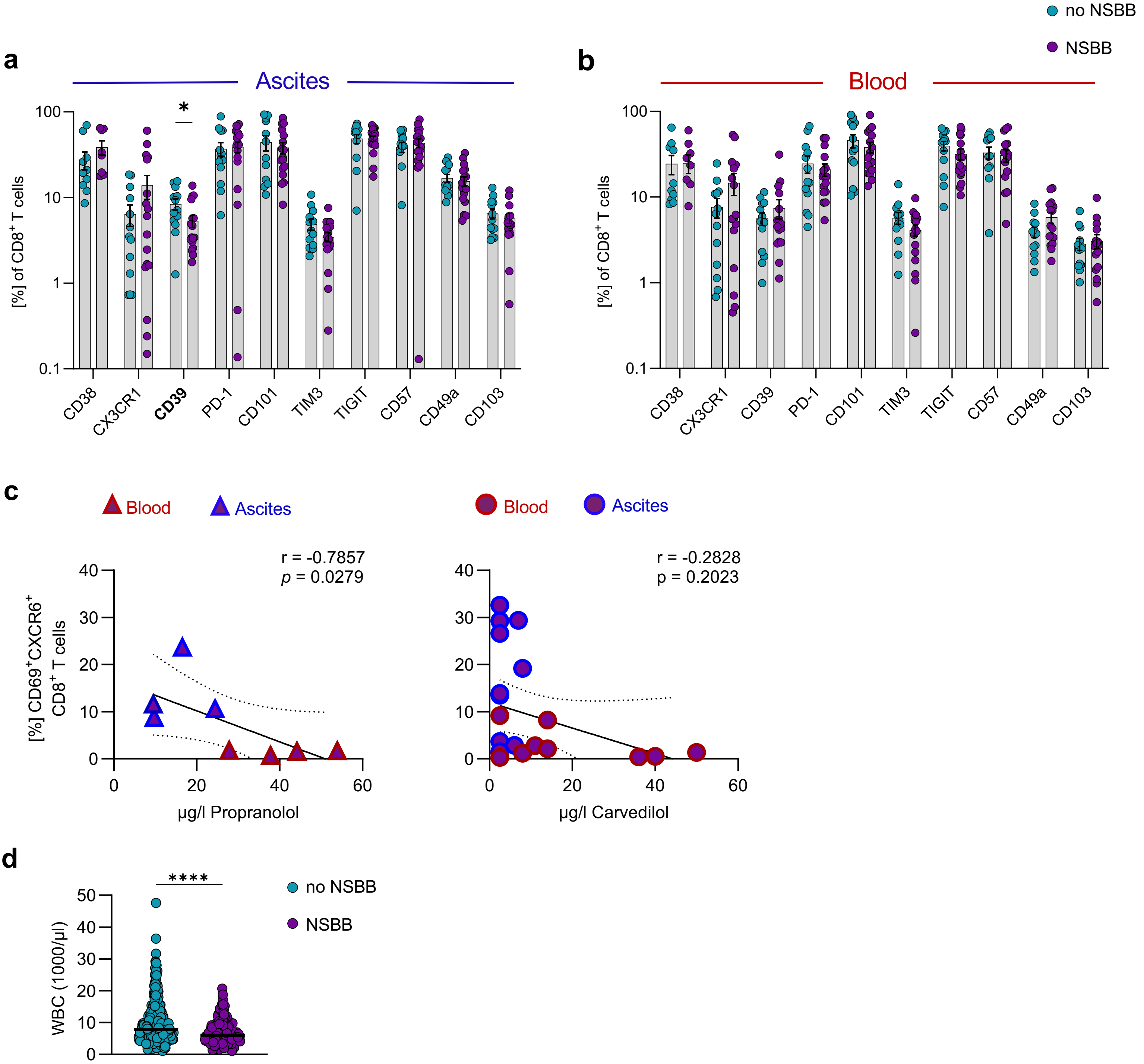

**Extended Data Table S1. Antibodies used for flow cytometric analyses.**

| Marker | Clone | Fluorophore | Company |
|---|---|---|---|
| ADRB1 | BSS-BS-0498R-PE | PE | Biozol |
| ADRB2 | orb15065 | FITC | biorbyt |
| CCR7 | G043H7 | PE/Fire 810 | BioLegend |
| CD101 | V7.1 | BUV563 | BD Horizon |
| CD103 | Ber-ACT8 | Brilliant Violet 711 | BioLegend |
| CD107a | H4A3 | Brilliant Violet 605 | BioLegend |
| CD127 | HIL-7R-M21 | APC-R700 | BD Horizon |
| CD14 | M5E2 | Alexa Fluor 700 | BD |
| CD14 | M5E2 | APC/Cyanine7 | BioLegend |
| CD16 | 3G8 | Brilliant Violet 510 | BioLegend |
| CD19 | HIB19 | Alexa Fluor 700 | BD |
| CD19 | SJ25C1 | APC/Cyanine8 | BioLegend |
| CD19 | SJ25C1 | APC/Fire 810 | BioLegend |
| CD3 | UCHT1 | APC/Fire 750 | BioLegend |
| CD3 | UCHT1 | BUV615 | BD Horizon |
| CD366 (Tim-3) | F38-2E2 | PE/Cyanine5 | BioLegend |
| CD38 | HB-7 | APC/Fire 810 | BioLegend |
| CD4 | RPA-T4 | PerCP/Cyanine5.5 | BioLegend |
| CD4 | SK3 | Brilliant Violet 786 | BD Horizon |
| CD4 | RPA-T4 | Brilliant Violet 570 | BioLegend |
| CD45 | HI30 | cFluor B548 | Cytek Biosciences |
| CD45RA | HI100 | Brilliant Violet 605 | BioLegend |
| CD49a | SR84 | BUV661 | BD Horizon |
| CD56 | 5.1H11 | PerCP/Cyanine5.5 | BioLegend |
| CD57 | QA17A04 | Brilliant Violet 510 | BioLegend |
| CD69 | FN50 | Brilliant Violet 421 | BioLegend |
| CD69 | FN50 | BUV737 | BD Horizon |
| CD8 | SK1 | Brilliant Violet 510 | BioLegend |
| CD8 | SK1 | Brilliant Violet 650 | BioLegend |
| CD8 | SK1 | BUV805 | BD Horizon |
| CX3CR1 | 2A9-1 | BUV496 | BD OptiBuild |
| CXCR6 | K041E5 | PE/Cyanine7 | BioLegend |

| | | | |
|---|---|---|---|
| CXCR6 | K041E5 | PE | BioLegend |
| EOMES | X4-83 | BUV395 | BD Horizon |
| Fixable Viability Stain 700 | | Alexa Fluor 700 | BD |
| Granzyme B | QA16A02 | PE/Dazzle 594 | BioLegend |
| Granzyme K | GM26E7 | APC | BioLegend |
| IFNγ | B27 | FITC | BioLegend |
| Ki-67 | SolA15 | Brilliant Violet 786 | Invitrogen |
| Ki-67 | SolA15 | Brilliant Violet 480 | Cytek Biosciences |
| LIVE/DEAD™ Fixable Blue Cell Stain Kit | | | Invitrogen |
| NKG2D | 1D11 | PE/Cyanine7 | BioLegend |
| PD-1 | EH12.2H7 | Brilliant Violet 650 | BioLegend |
| T-BET | 4B10 | Brilliant Violet 785 | BioLegend |
| TIGIT | MBSA43 | PerCP-eFluor 710 | eBioscience |
| TNFα | MAb11 | Brilliant Violet 650 | BioLegend |

**Extended Data Table S2. Reagents used in this study.**

| **Reagent** | **Company** |
|---|---|
| BD Cytofix/Cytoperm™ Fixation/Permeabilization Kit | BD |
| Brefeldin A | Sigma-Aldrich |
| CMV pp65 peptide | ProImmune |
| Dimethyl sulfoxide (DMSO) | Sigma-Aldrich |
| Fetal Calf Serum (FCS, heat-inactivated at 56 °C, 30 minutes) | Gibco |
| Foxp3/Transcription Factor Staining Buffer Set | eBioscience |
| Hanks' balanced salt solution (HBSS) | Gibco |
| HEPES buffer | Gibco |
| Human AB serum (heat-inactivate at 56 °C, 30 minutes) | PAN Biotech |
| Human IL-12 | Miltenyi Biotec |
| Human IL-15 | Miltenyi Biotec |
| Human IL-18 | MBL International Corporation |
| Ionomycin | Sigma-Aldrich |
| Isoproterenol hydrochloride (I6504) | Sigma-Aldrich |
| L-Glutamine | Gibco |
| Non-essential amino acids (NEAA) | Gibco |

| | |
|---|---|
| Penicillin/Streptomycin | Biochrom |
| Phorbol-12-myristate-13-acetate (PMA) | Sigma-Aldrich |
| Propranolol hydrochloride (P0884) | Sigma-Aldrich |
| purified anti-human CD28 | BioLegend |
| purified anti-human CD3 | BioLegend |
| RPMI-1640 with GlutaMAX ® | Gibco |
| Sodium azide solution (NaN3) | Merck |
| Sodium pyruvate | Gibco |
| Trypan blue solution | Gibco |

**Extended Data Table S3. Buffer and media used in this study.**

| **Buffer** | **Composition** |
|---|---|
| Erythrocyte lysis buffer | sterile water (H2O) |
| | 155 mM ammonium chloride (NH4Cl) |
| | 10 mM potassium hydrogen carbonate (KHCO3) |
| | 1 mM ethylenediaminetetraacetic acid (EDTA) |
| FACS buffer | Dulbecco's Phosphate-Buffered Saline (DPBS) |
| | 2 % FCS (heat-inactivate at 56 °C, 30 minutes) |
| | 0.05 % sodium azide solution (NaN3) |
| MACS buffer | Dulbecco's Phosphate-Buffered Saline (DPBS) |
| | 0.5 % BSA |
| | 2 mM EDTA |
| AB-Medium | RPMI-1640 with GlutaMAX ® |
| | 10 % human AB serum (heat-inactivated at 56 °C, 30 minutes) |
| | 1 % non-essential amino acids (NEAA) |
| | 1 % sodium pyruvate |
| | 1 % penicillin/streptomycin |
| | 5 mM HEPES buffer |
| | 1 mM L-glutamine |
| Freezing medium | 60 % FCS (heat-inactivate at 56 °C, 30 minutes) |
| | 30 % RPMI-1640 with GlutaMAX ® |
| | 10 % dimethyl sulfoxide (DMSO) |
